## Supplementary Material for "Joint inference of exclusivity patterns and recurrent trajectories from tumor mutation trees"

Xiang Ge Luo[1,2], Jack Kuipers[1,2], Niko Beerenwinkel[1,2*]
[1]Department of Biosystems Science and Engineering, ETH Zurich, Mattenstrasse 26, 4058 Basel, Switzerland
[2]SIB Swiss Institute of Bioinformatics, Mattenstrasse 26, 4058 Basel

**References 38**

### A Tree generating process and parameter estimation methods for TreeMHN

#### A.1 Tree generating process pseudocode

In Algorithm 1 we provide the pseudocode for the tree-generating process described in the Methods section. A tree $\mathscr{T}$ consists of a vertex set $V$ and an edge set $E$. Each node in the vertex set is of the form $(i, \pi_i)$, where $i$ is the unique index of the node in $\mathscr{T}$, and $\pi_i$ is a mutational trajectory that runs from the root to the node $i$. An illustrative example is given in Supplementary Figure S1(a) and (b).

**Algorithm 1** Tree generating process

**Input**
- $n$ — Number of mutations
- $\Theta$ — Mutual Hazard Network represented by an $n$-by-$n$ matrix
- $\lambda_s$ — Sampling rate

**Output**
- $\mathscr{T}$ — A tree structure

$t_s \sim \text{Exp}(\lambda_s)$ ▷ Draw a sampling time
$V \leftarrow \{(1, (0))\}$ ▷ The root node has index 1
$E \leftarrow \emptyset$ ▷ The edge set is empty
$t_{(0)} = 0$ ▷ The root node is at time 0
$\mathcal{U}_{\text{current}} \leftarrow \{1\}$ ▷ Index set of the nodes to be visited
**while** $\mathcal{U}_{\text{current}}$ is non-empty **do**
  $\mathcal{U}_{\text{next}} \leftarrow \{\}$ ▷ Initialize the set of nodes to visit next
  **for** $i$ in $\mathcal{U}_{\text{current}}$ **do**
    **for** $j$ in $[n] \setminus \pi_i$ **do**
      $\pi \leftarrow (\pi_i, j)$ ▷ Extend the trajectory $\pi_i$ by one mutation in $[n] \setminus \pi_i$
      $t_\pi \sim t_{\pi_i} + \text{Exp}(\lambda_{(\pi_i,j)})$, $\lambda_{(\pi_i,j)} = \Theta_{jj} \prod_{l \in \pi_i} \Theta_{jl}$ ▷ Draw a waiting time for $\pi$
      **if** $t_\pi < t_s$ **then** ▷ Expand the tree by one node
        $k \leftarrow |V| + 1$
        $V \leftarrow V \cup \{(k, \pi)\}$
        $E \leftarrow E \cup \{i \rightarrow k\}$
        $\mathcal{U}_{\text{next}} \leftarrow \mathcal{U}_{\text{next}} \cup \{k\}$
      **end if**
    **end for**
  **end for**
  $\mathcal{U}_{\text{current}} \leftarrow \mathcal{U}_{\text{next}}$
**end while**
$\mathscr{T} \leftarrow (V, E)$

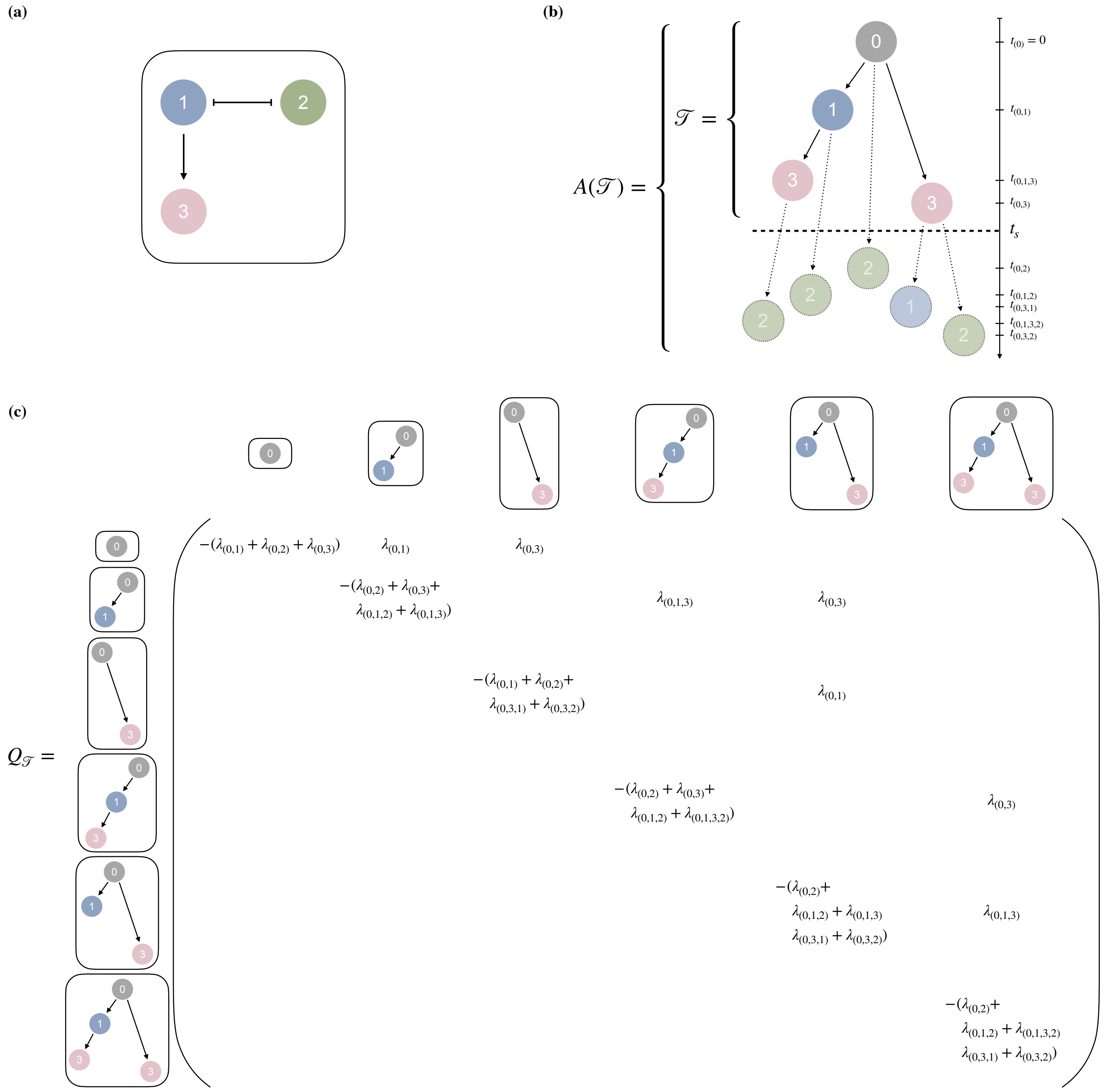

Figure S1: (a) A Mutual Hazard Network with three distinct mutations. (b) An example of a tree generated from the MHN in (a), where both the observed part $\mathcal{T}$ and the augmented tree $A(\mathcal{T})$ are shown. (c) The sub-matrix $Q_{\mathcal{T}}$, which corresponds to the tree $\mathcal{T}$ in (b) and is defined in Supplementary Eq. (12).

#### A.2 Tree generating process as a continuous-time Markov chain

In this section, we provide an alternative definition of the tree generating process in terms of a continuous-time Markov chain on the state space of tumor mutation trees. Recall that a tumor mutation tree $\mathcal{T}$ is a rooted tree representing the evolutionary history of a tumor. Each vertex $v$ in $\mathcal{T}$ corresponds to a subclone $\pi$, uniquely defined by the sequence of mutations along the trajectory from the root to $v$. Mathematically, we write $\pi = (0, \sigma_1, \ldots, \sigma_d)$, where $d \in \{0, \ldots, n\}$ is the length of the trajectory, $n$ is the total number of mutations being considered in the model, and $\sigma_i$ are non-duplicated elements in $[n] = \{1, \ldots, n\}$. The root is a special subclone with no mutations, denoted by $(0)$. We denote the parent of a subclone $\pi = (0, \sigma_1, \ldots, \sigma_d)$ by $\text{pa}(\pi) = (0, \sigma_1, \ldots, \sigma_{d-1})$ and its children by $\text{ch}(\pi) = \{\pi' = (0, \sigma_1, \ldots, \sigma_d, \sigma_{d+1}) \mid \sigma_{d+1} \in [n] \setminus \{\sigma_1, \ldots, \sigma_d\}\}$.

All subclones except the root in $\mathscr{T}$ satisfy the following partial order relation: if $\pi$ is a subclone of $\mathscr{T}$, then its parent $\text{pa}(\pi)$ must also be a subclone of $\mathscr{T}$ with an edge $\text{pa}(\pi) \to \pi$, since by definition $\pi$ can only occur after $\text{pa}(\pi)$. We can view a tree $\mathscr{T}$ as a set of subclones compatible with the partial order relation and omit the implicitly implied edge set. For example, the tree displayed in Supplementary Figure S1(b) consists of four subclones $\{(0), (0,1), (0,3), (0,1,3)\}$, and one can read off the edges by enumerating all parent-child pairs, *i.e.* $(0) \to (0,1)$, $(0) \to (0,3)$, and $(0,1) \to (0,1,3)$. Removing a leaf like $(0,1,3)$ from the set results in another smaller tree with three vertices.

Since tumor mutation trees are effectively sets of subclones, the notations and binary operations on sets can be applied directly:

- $\pi \in \mathscr{T}$ means that $\pi$ is a subclone of the tree $\mathscr{T}$.
- We use $|\mathscr{T}|$ to denote the number of subclones in tree $\mathscr{T}$.
- We say that $\mathscr{T}_1$ is a subtree of $\mathscr{T}_2$ if all subclones in $\mathscr{T}_1$ can be found in $\mathscr{T}_2$ and write $\mathscr{T}_1 \subseteq \mathscr{T}_2$. If $\mathscr{T}_1$ is a subtree of $\mathscr{T}_2$ and $|\mathscr{T}_1| < |\mathscr{T}_2|$, then we write $\mathscr{T}_1 \subset \mathscr{T}_2$.
- We use $\mathscr{T}_2 \setminus \mathscr{T}_1$ to denote the set of subclones in $\mathscr{T}_2$ that are not subclones of $\mathscr{T}_1$. The number of such subclones are denoted by $|\mathscr{T}_2 \setminus \mathscr{T}_1|$.
- We use $\mathscr{T}_1 \cup \mathscr{T}_2$ to denote the union of subclones in $\mathscr{T}_1$ and $\mathscr{T}_2$, which constitutes a valid tree since the partial order relation still holds. We may also use $\mathscr{T} \cup S$ to denote the union of subclones in a tree $\mathscr{T}$ and a set of subclones $S$, which may not satisfy the partial order relation.
- Given a tree $\mathscr{T}$, we define the exit set of $\mathscr{T}$ as

  $$\text{Exit}(\mathscr{T}) = \{\pi \mid \pi \notin \mathscr{T}, \text{pa}(\pi) \in \mathscr{T}\},$$

  which contains the children of the existing subclones that are not yet in $\mathscr{T}$, *i.e.* all subclones that could appear next. Then, we define the augmented tree as

  $$A(\mathscr{T}) := \mathscr{T} \cup \text{Exit}(\mathscr{T}), \quad \textit{i.e.}, \ \text{Exit}(\mathscr{T}) = A(\mathscr{T}) \setminus \mathscr{T}.$$

We model tumor progression as a continuous-time Markov chain $(X_t)_{t \geq 0}$ [1] on the state space of all tumor mutation trees, denoted by $S^n_{\mathscr{T}}$. The size of the state space grows super-exponentially in the number of mutations $n$ since

$$|S^1_{\mathscr{T}}| = 2, \qquad |S^n_{\mathscr{T}}| = \left(|S^{n-1}_{\mathscr{T}}| + 1\right)^n. \tag{1}$$

The initial state of the chain is the tree with only the root, $X_0 = \mathscr{T}_0$, meaning that no mutations of interest are present at the start of the evolution. It is equivalent to say that the initial distribution of the chain has a point mass of 1 on $\mathscr{T}_0$ and 0 on all other trees at time $t = 0$, denoted by the Kronecker delta $\delta_{\mathscr{T}_0}$. The generator $Q$-matrix $Q = (q_{\mathscr{T}_1,\mathscr{T}_2} : \mathscr{T}_1, \mathscr{T}_2 \in S^n_{\mathscr{T}})$ is indexed by trees and defined as

$$q_{\mathscr{T}_1,\mathscr{T}_2} := \begin{cases} \lambda_{\mathscr{T}_2 \setminus \mathscr{T}_1} & \mathscr{T}_1 \subset \mathscr{T}_2, |\mathscr{T}_2 \setminus \mathscr{T}_1| = 1 \\ -\sum_{\mathscr{T} \neq \mathscr{T}_1} q_{\mathscr{T}_1,\mathscr{T}} & \mathscr{T}_1 = \mathscr{T}_2 \\ 0 & \text{otherwise.} \end{cases} \tag{2}$$

The dimension of $Q$ is $|S^n_{\mathscr{T}}| \times |S^n_{\mathscr{T}}|$. We say $(X_t)_{t \geq 0}$ is continuous-time *Markov*$(\delta_{\mathscr{T}_0}, Q)$. The interpretation of the off-diagonal entries is that transitioning from $\mathscr{T}_1$ to $\mathscr{T}_2$ is only possible if $\mathscr{T}_2$ has exactly one extra subclone than $\mathscr{T}_1$, and this new subclone $\pi = \mathscr{T}_2 \setminus \mathscr{T}_1$ must be a child of one of the existing subclones in $\mathscr{T}_1$, *i.e.* $\pi \in \text{Exit}(\mathscr{T}_1)$. The transition is achieved by accumulating one subclone at a time, and this process is irreversible. The transition rate is associated with all mutations in $\pi = (\text{pa}(\pi), i)$ for some $i \in [n] \setminus \text{pa}(\pi)$ and parameterized by a Mutual Hazard Network $\Theta \in \mathbb{R}^{n \times n}$,

$$\lambda_{\mathscr{T}_2 \setminus \mathscr{T}_1} = \lambda_\pi := \exp\left(\theta_{ii} + \sum_{j \in \text{pa}(\pi)} \theta_{ij}\right) = \Theta_{ii} \prod_{j \in \text{pa}(\pi)} \Theta_{ij}, \tag{3}$$

which is equivalent to the definition in Equation (1) of the main text (notation-wise shifted by one mutation). The diagonal entries are chosen such that the row sums of $Q$ are zero. We interpret the off-diagonal row sum

$$q_{\mathscr{T}_1} := \sum_{\mathscr{T} \neq \mathscr{T}_1} q_{\mathscr{T}_1,\mathscr{T}} = \sum_{\pi \in \mathrm{Exit}(\mathscr{T}_1)} \lambda_\pi,$$

as the rate of leaving the state $\mathscr{T}_1$, which is the sum of the rates of all events that could happen next.

Following Chapter 2 of [1], we define the jump times $J_0, J_1, \ldots$ and the holding times $S_1, S_2, \ldots$ of $(X_t)_{t\geq 0}$ to be

$$J_0 = 0, \quad J_{k+1} = \inf\{t \geq J_k : X_t \neq X_{J_k}\}$$

for $k = 0, 1, \ldots$, with $\inf \emptyset = \infty$, and for $k = 1, 2, \ldots$,

$$S_k = \begin{cases} J_k - J_{k-1} & J_{k-1} < \infty \\ \infty & \text{otherwise.} \end{cases}$$

Furthermore, we define the jump matrix $\Xi = (\xi_{\mathscr{T}_1,\mathscr{T}_2} : \mathscr{T}_1, \mathscr{T}_2 \in S^n_{\mathscr{T}})$ of $Q$ as

$$\begin{aligned} \xi_{\mathscr{T}_1,\mathscr{T}_2} &:= \begin{cases} q_{\mathscr{T}_1,\mathscr{T}_2}/q_{\mathscr{T}_1} & \mathscr{T}_1 \neq \mathscr{T}_2 \text{ and } q_{\mathscr{T}_1} \neq 0 \\ 0 & \mathscr{T}_1 \neq \mathscr{T}_2 \text{ and } q_{\mathscr{T}_1} = 0, \end{cases} \\ \xi_{\mathscr{T}_1,\mathscr{T}_1} &:= \begin{cases} 0 & q_{\mathscr{T}_1} \neq 0 \\ 1 & q_{\mathscr{T}_1} = 0. \end{cases} \end{aligned} \tag{4}$$

By Theorem 2.8.2 of [1], the jump chain $(Y_k)_{k\geq 0}$ associated with $(X_t)_{t\geq 0}$ is a discrete-time Markov chain with initial distribution $\delta_{\mathscr{T}_0}$ and jump matrix $\Xi$, where the entry $\xi_{\mathscr{T}_1,\mathscr{T}_2}$ is the probability of jumping from $\mathscr{T}_1$ to $\mathscr{T}_2$. We say $(Y_k)_{k\geq 0}$ is discrete-time *Markov*$(\delta_{\mathscr{T}_0}, \Xi)$. In particular, conditioned on the chain of states $Y_0, Y_1, \ldots, Y_{k-1}$ for each $k \geq 1$, the holding times $S_1, S_2, \ldots, S_k$ are independent exponential random variables with parameters $q_{Y_0}, q_{Y_1}, \ldots, q_{Y_{k-1}}$.

Next we show the equivalence between the formulation in terms of Markov chains above and the tree-generating process defined in the main text.

**Theorem 1.** *A right-continuous process $(X_t)_{t\geq 0}$ with values in the space of tumor mutation trees $S^n_{\mathscr{T}}$ is continuous-time Markov$(\delta_{\mathscr{T}_0}, Q)$ if and only if the waiting times of subclones $\pi \in \mathscr{T}$ for any $\mathscr{T} \in S^n_{\mathscr{T}}$ are exponential random variables such that*

$$T_{(0)} = 0, \qquad T_\pi \sim T_{\mathrm{pa}(\pi)} + \mathrm{Exp}(\lambda_\pi). \tag{5}$$

*Proof.* ($\Longrightarrow$) Suppose $(X_t)_{t\geq 0}$ is continuous-time *Markov*$(\delta_{\mathscr{T}_0}, Q)$. The initial state $X_0 = \mathscr{T}_0$ corresponds to the initial wild-type subclone (0) at time 0, *i.e.* $T_{(0)} = 0$. By the infinitesimal definition in Theorem 2.8.2 of [1], for all $t, h \geq 0$, conditioned on $X_t = \mathscr{T}_1$ for some $\mathscr{T}_1 \in S^n_{\mathscr{T}}$, $X_{t+h}$ is independent of $(X_s : s \leq t)$. As $h \downarrow 0$, uniformly in $t$, for all $\mathscr{T}_2 \in S^n_{\mathscr{T}}$,

$$P(X_{t+h} = \mathscr{T}_2 \mid X_t = \mathscr{T}_1) = \delta_{\mathscr{T}_1,\mathscr{T}_2} + q_{\mathscr{T}_1,\mathscr{T}_2} h + o(h),$$

where the Kronecker delta $\delta_{\mathscr{T}_1,\mathscr{T}_2} = 1$ if $\mathscr{T}_1 = \mathscr{T}_2$ and 0 otherwise. For any subclone $\pi \neq (0)$, we can construct a smaller chain $(Z^\pi_t)_{t\geq 0}$ on two states $\{\mathrm{pa}(\pi), \pi\}$ with initial state $Z^\pi_0 = \mathrm{pa}(\pi)$ from $(X_t)_{t\geq 0}$, in order to obtain the marginal probability distribution of $\pi$. For all $t \geq 0$, as $h \to 0$,

$$\begin{aligned} P(Z^\pi_{t+h} = \pi, Z^\pi_t = \mathrm{pa}(\pi)) &= \sum_{\mathscr{T}:\mathrm{pa}(\pi)\in\mathscr{T},\pi\notin\mathscr{T}} P(X_{t+h} = \mathscr{T} \cup \{\pi\}, X_t = \mathscr{T}) \\ &= \sum_{\mathscr{T}:\mathrm{pa}(\pi)\in\mathscr{T},\pi\notin\mathscr{T}} P(X_{t+h} = \mathscr{T} \cup \{\pi\} \mid X_t = \mathscr{T}) P(X_t = \mathscr{T}) \\ &= \sum_{\mathscr{T}:\mathrm{pa}(\pi)\in\mathscr{T},\pi\notin\mathscr{T}} (\lambda_\pi + o(h)) P(X_t = \mathscr{T}) \\ &= (\lambda_\pi + o(h)) \sum_{\mathscr{T}:\mathrm{pa}(\pi)\in\mathscr{T},\pi\notin\mathscr{T}} P(X_t = \mathscr{T}) \\ &= (\lambda_\pi + o(h)) P(Z^\pi_t = \mathrm{pa}(\pi)). \end{aligned}$$

Hence,

$$\begin{aligned}
P(Z^\pi_{t+h} = \pi \mid Z^\pi_t = \mathrm{pa}(\pi)) &= \frac{P(Z^\pi_{t+h} = \pi, Z^\pi_t = \mathrm{pa}(\pi))}{P(Z_t = \mathrm{pa}(\pi))} \\
&= \frac{(\lambda_\pi + o(h))P(Z^\pi_t = \mathrm{pa}(\pi))}{P(Z^\pi_t = \mathrm{pa}(\pi))} \\
&= \lambda_\pi + o(h),
\end{aligned}$$

$$P(Z^\pi_{t+h} = \mathrm{pa}(\pi) \mid Z^\pi_t = \mathrm{pa}(\pi)) = 1 - \lambda_\pi + o(h).$$

Similarly,

$$P(Z^\pi_{t+h} = \mathrm{pa}(\pi) \mid Z^\pi_t = \pi) = o(h), \quad \text{and} \quad P(Z^\pi_{t+h} = \pi \mid Z^\pi_t = \pi) = 1 + o(h).$$

By Theorem 2.8.2 of [1], $(Z^\pi_t)_{t\geq 0}$ is also a continuous-time Markov chain with the generator matrix

$$\begin{array}{c} \\ \mathrm{pa}(\pi) \\ \pi \end{array} \begin{array}{c} \begin{array}{cc} \mathrm{pa}(\pi) & \pi \end{array} \\ \begin{bmatrix} -\lambda_\pi & \lambda_\pi \\ 0 & 0 \end{bmatrix} \end{array}.$$

It follows that the holding time from $\mathrm{pa}(\pi)$ to $\pi$ is an exponential random variable with rate $\lambda_\pi$, as required.

($\Longleftarrow$) Let $(X_t)_{t\geq 0}$ be a right-continuous process with values in $S^n_{\mathscr{T}}$, which tracks the change of state resulting from the occurrence of new subclones with exponentially-distributed waiting times as defined in Supplementary Eq. (5). Let $(Y_k)_{k\geq 0}$ be the associated jump chain. We will use the jump chain/holding time definition in Theorem 2.8.2 of [1] to prove that $(X_t)_{t\geq 0}$ is continuous-time *Markov*$(\delta_{\mathscr{T}_0}, Q)$.

At time $t = 0$, there is only the initial wild-type subclone $\pi = (0)$, which corresponds to the initial state $X_0 = Y_0 = \mathscr{T}_0$. Let $W(k)$ be the statement that for each $k \geq 1$, conditioned on the chain of states $Y_0, Y_1, \ldots, Y_{k-1}$, the holding times $S_1, S_2, \ldots, S_k$ are independent exponential random variables with parameters $q_{Y_0}, q_{Y_1}, \ldots, q_{Y_{k-1}}$. We will prove $W(k)$ by induction on $k \geq 1$.

- Base case: Conditioned on $Y_0 = \mathscr{T}_0$, we have

$$S_1 = J_1 = \inf\{t \geq 0 : X_t \neq \mathscr{T}_0\} = \inf\{T_{(0,i)} \geq 0 : i \in [n]\},$$

because the set of mutations that could happen next is $[n]$, and whichever mutation appears first will change the state of the chain. Each waiting time $T_{(0,i)}$ is an independent exponential random variable with rate $\lambda_{(0,i)}$. By Theorem 2.3.3 of [1] (competing exponentials), the first holding time $S_1$ is an exponential random variable with rate

$$\sum_{i\in[n]} \lambda_{(0,i)} = \sum_{\pi\in \mathrm{Exit}(\mathscr{T}_0)} \lambda_\pi = q_{\mathscr{T}_0} = q_{Y_0},$$

independent of the next state $Y_1$, which has distribution $(\xi_{\mathscr{T}_0,\mathscr{T}_1} : \mathscr{T}_1 \in S^n_{\mathscr{T}})$ because

$$P(Y_1 = \mathscr{T}_1 \mid Y_0 = \mathscr{T}_0) = \begin{cases} \frac{\lambda_{(0,i)}}{\sum_{i\in[n]} \lambda_{(0,i)}} = \frac{q_{\mathscr{T}_0,\mathscr{T}_1}}{q_{\mathscr{T}_0}} & \mathscr{T}_1 = \mathscr{T}_0 \cup \{(0,i)\} \text{ for } i \in [n] \\ 0 & \text{otherwise.} \end{cases}$$

Hence, the statement $W(1)$ is true.

- Induction step: Assume that for a given value $k = l \geq 1$, $W(k)$ holds. Next, conditioned on $Y_l = \mathscr{T}_l \in S^n_{\mathscr{T}}$, the next holding time depends on the waiting times of the subclones that can appear next,

$$S_{l+1} = J_{l+1} - J_l = \inf\{t \geq J_l : X_t \neq X_{J_l}\} - J_l = \inf\{T_\pi - J_l \geq 0 : \pi \in \mathrm{Exit}(\mathscr{T}_l)\}.$$

We first assume that $\mathrm{Exit}(\mathscr{T}_l) \neq \emptyset$. For each $\pi \in \mathrm{Exit}(\mathscr{T}_l)$, by definition, we have $\mathrm{pa}(\pi) \in \mathscr{T}_l$ and $T_{\mathrm{pa}(\pi)} \leq J_l \Leftrightarrow J_l - T_{\mathrm{pa}(\pi)} \geq 0$. By Supplementary Eq. (5), the waiting time from $\mathrm{pa}(\pi)$ to $\pi$ is an exponentially-distributed random variable,

$$T_\pi - T_{\mathrm{pa}(\pi)} \sim \mathrm{Exp}(\lambda_\pi),$$

then by Theorem 2.3.1 of [1] (memoryless property), the waiting time from the most recent jump to the occurrence of $\pi$ remains exponentially distributed with the same rate,

$$T_\pi - J_l = (T_\pi - T_{\mathrm{pa}(\pi)}) - (J_l - T_{\mathrm{pa}(\pi)}) \sim \mathrm{Exp}(\lambda_\pi),$$

and independent of $T_{\pi'} - J_l$ for $\pi' \neq \pi \in \mathscr{T}_l$. By Theorem 2.3.3 of [1], $S_{l+1}$ is an exponential random variable with rate

$$\sum_{\pi \in \mathrm{Exit}(Y_l)} \lambda_\pi = q_{\mathscr{T}_l} = q_{Y_l},$$

independent of $S_1, S_2, \ldots, S_l$ as required. $S_{l+1}$ is also independent of the next state $Y_{l+1}$, which has distribution $(\xi_{\mathscr{T}_l, \mathscr{T}_{l+1}} : \mathscr{T}_{l+1} \in S^n_{\mathscr{T}})$ because

$$P(Y_{l+1} = \mathscr{T}_{l+1} \mid Y_l = \mathscr{T}_l) = \begin{cases} \frac{\lambda_\pi}{\sum_{\pi' \in \mathrm{Exit}(Y_l)} \lambda_{\pi'}} = \frac{q_{\mathscr{T}_l, \mathscr{T}_{l+1}}}{q_{\mathscr{T}_l}} & \mathscr{T}_{l+1} = \mathscr{T}_l \cup \{\pi\} \text{ for } \pi \in \mathrm{Exit}(Y_l) \\ 0 & \text{otherwise.} \end{cases}$$

If $\mathrm{Exit}(\mathscr{T}_l) = \emptyset$, *i.e.* no subclones can appear next, then by the definition in [1], $S_{l+1} = \inf \emptyset = \infty$ is exponentially distributed with rate $q_{\mathscr{T}_l} = 0$, and $Y_{l+1} = Y_l = \mathscr{T}_l$ is an absorbing state with $P(Y_{l+1} = \mathscr{T}_l \mid Y_l = \mathscr{T}_l) = 1$.

Therefore, the statement $W(l+1)$ is also true. By induction, $W(k)$ holds for all $k \geq 1$. Since $Y_0 = \mathscr{T}_0$ and the transition probabilities $P(Y_k = \mathscr{T}_{k-1} \mid Y_{k-1} = \mathscr{T}_{k-1})$ for $k \geq 1$ match the definition of $\Xi$ in Supplementary Eq. (4), the jump chain $(Y_k)_{k \geq 0}$ is discrete-time *Markov*$(\delta_{\mathscr{T}_0}, \Xi)$. By the jump chain/holding time definition in Theorem 2.8.2 of [1], $(X_t)_{t \geq 0}$ is continuous-time *Markov*$(\delta_{\mathscr{T}_0}, Q)$.

We have proved that $(X_t)_{t \geq 0}$ is continuous-time *Markov*$(\delta_{\mathscr{T}_0}, Q)$ if and only if the subclones $\pi \in \mathscr{T}$ for any $\mathscr{T} \in S^n_{\mathscr{T}}$ have exponentially-distributed waiting times as defined in Supplementary Eq. (5). □

**Theorem 2.** *Let $\Theta \in \mathbb{R}^{n \times n}$ be a Mutual Hazard Network, where $n \in \mathbb{N}^+$ is the number of mutations. Let $(X_t)_{t \geq 0}$ be continuous-time Markov$(\delta_{\mathscr{T}_0}, Q)$. Suppose $\mathscr{T} \in S^n_{\mathscr{T}}$ is an output of the tree generating process. Then the marginal probability of observing $\mathscr{T}$ is given by*

$$p(\mathscr{T} \mid \Theta) := P\left( \max_{\pi \in \mathscr{T}} T_\pi < T_s < \min_{\pi' \in \mathrm{Exit}(\mathscr{T})} T_{\pi'} \;\middle|\; \Theta \right) = \left( \lambda_s (\lambda_s I - Q)^{-1} \right)_{\mathscr{T}_0, \mathscr{T}}. \tag{6}$$

*Proof.* In the tree generating process, the waiting times of the subclones are exponential random variables as defined in Supplementary Eq. (5). By Theorem 1, the right-continuous process $(X_t)_{t \geq 0}$ that tracks the change of state resulting from the occurrence of new subclones is continuous-time *Markov*$(\delta_{\mathscr{T}_0}, Q)$. That is, for all $t > 0$, and $\mathscr{T} \in S^n_{\mathscr{T}}$,

$$X_t = \mathscr{T} \quad \Leftrightarrow \quad \max_{\pi \in \mathscr{T}} T_\pi < t < \min_{\pi' \in \mathrm{Exit}(\mathscr{T})} T_{\pi'},$$

because if there exists $\pi \in \mathscr{T}$ such that $T_\pi > t$, or $\pi' \in \mathrm{Exit}(\mathscr{T})$ such that $T_{\pi'} < t$, then $X_t \neq \mathscr{T}$, which is a contradiction. Next, the matrix $P(t) = e^{tQ}$ is the transition probability matrix of $(X_t)_{t \geq 0}$, which by Theorem 2.1.1 of [1] solves the backward equation,

$$\frac{d}{dt} P(t) = QP(t), \qquad P(0) = I. \tag{7}$$

The transition probability from $\mathscr{T}_1$ to $\mathscr{T}_2$ in time $t$ is given by

$$P(X_t = \mathscr{T}_2 \mid X_0 = \mathscr{T}_1) = p_{\mathscr{T}_1, \mathscr{T}_2}(t),$$

where $p_{\mathscr{T}_1, \mathscr{T}_2}(t)$ is the $(\mathscr{T}_1, \mathscr{T}_2)$ entry in $e^{tQ}$, the row is indexed by $\mathscr{T}_1$, and the column is indexed by $\mathscr{T}_2$. In particular, starting with $X_0 = \mathscr{T}_0$, the probability of observing $\mathscr{T}$ at sampling time $t_s$ is

$$P(X_{t_s} = \mathscr{T} \mid X_0 = \mathscr{T}_0) = p_{\mathscr{T}_0, \mathscr{T}}(t_s) = \left( e^{t_s Q} \right)_{\mathscr{T}_0, \mathscr{T}}, \tag{8}$$

which is the $(\mathscr{T}_0, \mathscr{T})$ entry in $e^{t_s Q}$. It follows that

$$P\left(\max_{\pi \in \mathscr{T}} T_\pi < t_s < \min_{\pi' \in \text{Exit}(\mathscr{T})} T_{\pi'} \;\middle|\; \Theta\right) = P(X_{t_s} = \mathscr{T} \mid X_0 = \mathscr{T}_0) = \left(e^{t_s Q}\right)_{\mathscr{T}_0, \mathscr{T}}. \tag{9}$$

In the tree generating process, we assume that the unknown sampling time follows an independent exponential distribution $T_s \sim \text{Exp}(\lambda_s)$ with density function $f_{\lambda_s}(t_s) = \lambda_s \exp(-\lambda_s t_s)$. Integrating it out gives us

$$\begin{aligned} P\left(\max_{\pi \in \mathscr{T}} T_\pi < T_s < \min_{\pi' \in \text{Exit}(\mathscr{T})} T_{\pi'} \;\middle|\; \Theta\right) &= \int_0^\infty f_{\lambda_s}(t_s) P\left(\max_{\pi \in \mathscr{T}} T_\pi < t_s < \min_{\pi' \in \text{Exit}(\mathscr{T})} T_{\pi'} \;\middle|\; \Theta\right) dt_s \\ &= \left(\int_0^\infty f_{\lambda_s}(t_s) e^{t_s Q} dt_s\right)_{\mathscr{T}_0, \mathscr{T}} \\ &= \left(\int_0^\infty \lambda_s e^{-\lambda_s t_s} e^{t_s Q} dt_s\right)_{\mathscr{T}_0, \mathscr{T}} \\ &= \left(\lambda_s \int_0^\infty e^{-\lambda_s t_s} e^{t_s Q} dt_s\right)_{\mathscr{T}_0, \mathscr{T}}. \end{aligned} \tag{10}$$

Here, $\mathcal{L}(P(t)) = \int_0^\infty e^{-\lambda_s t_s} P(t_s) dt_s = \int_0^\infty e^{-\lambda_s t_s} e^{t_s Q} dt_s = A(\lambda_s)$ is the Laplace transform of $P(t)$ [2]. By the Laplace transform of derivatives [2], we have

$$\mathcal{L}\left(\frac{d}{dt} P(t)\right) = \lambda_s A(\lambda_s) - P(0) = \lambda_s A(\lambda_s) - I.$$

Applying Laplace transforms to both sides of Supplementary Eq. (7) gives

$$\lambda_s A(\lambda_s) - I = Q A(\lambda_s) \quad \Leftrightarrow \quad (\lambda_s I - Q) A(\lambda_s) = I.$$

By the definition in Supplementary Eq. (2), we can permute the rows and columns of the generator matrix $Q$ such that $Q$ is upper-triangular, *e.g.* by sorting the trees by increasing tree size, such that the matrix $V := \lambda_s I - Q$ is also upper-triangular. Since all transition rates are positive by Supplementary Eq. (3), the diagonal entries of $Q$ are non-positive. Hence, the diagonal entries of $V$ are positive. It follows that the determinant of $V$, as the product of the diagonal entries, is non-zero, *i.e.* $\det(V) \neq 0$. Hence, $V$ is non-singular, $V^{-1}$ exists, and

$$A(\lambda_s) = (\lambda_s I - Q)^{-1}.$$

Therefore, the marginal probability of observing $\mathscr{T}$ is given by

$$p(\mathscr{T} \mid \Theta) := P\left(\max_{\pi \in \mathscr{T}} T_\pi < T_s < \min_{\pi' \in \text{Exit}(\mathscr{T})} T_{\pi'} \;\middle|\; \Theta\right) = (\lambda_s A(\lambda_s))_{\mathscr{T}_0, \mathscr{T}} = \left(\lambda_s (\lambda_s I - Q)^{-1}\right)_{\mathscr{T}_0, \mathscr{T}}.$$

□

#### A.3 Maximum likelihood estimation of an MHN given a set of trees

In the previous section, we have shown in Theorem 2 that computing $p(\mathscr{T} \mid \Theta)$ involves inverting the matrix $V = \lambda_s I - Q$ of dimension $|S^n_{\mathscr{T}}| \times |S^n_{\mathscr{T}}|$, where $|S^n_{\mathscr{T}}|$ grows super-exponentially in $n$ (Supplementary Eq. (1)). Hence, the matrix inversion is intractable for even a moderate size $n$. In this section, we seek to simplify the calculation of Supplementary Eq. (6).

We sort the rows and columns of $Q$ by increasing tree size $(\mathscr{T}_0, \mathscr{T}_1, \dots \mathscr{T}_L)$ such that $|\mathscr{T}_0| < |\mathscr{T}_1| < \cdots < |\mathscr{T}_L|$, where $L := |S^n_{\mathscr{T}}|$ is the total number of trees in the state space. In other words, $Q$ is upper-triangular, and so is $V$. Let $\boldsymbol{e}_{\mathscr{T}}$ be a unit vector on $S^n_{\mathscr{T}}$ with 1 at the position of $\mathscr{T} \in S^n_{\mathscr{T}}$ and 0 at other positions. The dimension of $\boldsymbol{e}_{\mathscr{T}}$ is $L$. For example, $\boldsymbol{e}_{\mathscr{T}_0} = (1, 0, \dots, 0)^\top$. Then, we can rewrite Supplementary Eq. (6) as

$$p(\mathscr{T} \mid \Theta) = \left(\lambda_s V^{-1}\right)_{\mathscr{T}_0, \mathscr{T}} = \lambda_s \boldsymbol{e}_{\mathscr{T}_0}^\top V^{-1} \boldsymbol{e}_{\mathscr{T}}.$$

To compute $\boldsymbol{x}^\top := \boldsymbol{e}_{\mathscr{T}_0}^\top V^{-1}$, we can solve the equation $V^\top \boldsymbol{x} = \boldsymbol{e}_{\mathscr{T}_0}$ with forward substitution because $V^\top$ is lower-triangular:

$$\begin{array}{ccccccccc}
V_{\mathscr{T}_0,\mathscr{T}_0} x_0 & & & & & & & = & 1 \\
V_{\mathscr{T}_0,\mathscr{T}_1} x_0 & + & V_{\mathscr{T}_1,\mathscr{T}_1} x_1 & & & & & = & 0 \\
\vdots & & \vdots & & \ddots & & & & \vdots \\
V_{\mathscr{T}_0,\mathscr{T}_L} x_0 & + & V_{\mathscr{T}_1,\mathscr{T}_L} x_1 & + & \dots & + & V_{\mathscr{T}_L,\mathscr{T}_L} x_L & = & 0
\end{array}$$

The recursive formula is

$$\begin{aligned}
x_0 &= V^{-1}_{\mathscr{T}_0,\mathscr{T}_0} \\
x_m &= \frac{-\sum_{l=0}^{m-1} V_{\mathscr{T}_l,\mathscr{T}_m} x_l}{V_{\mathscr{T}_m,\mathscr{T}_m}}, \qquad m = 1, 2, \dots, L.
\end{aligned} \tag{11}$$

To compute $p(\mathscr{T} \mid \Theta)$, it is unnecessary to compute the entire vector $\boldsymbol{x}$, because $\boldsymbol{e}_{\mathscr{T}_0}^\top V^{-1} \boldsymbol{e}_{\mathscr{T}} = \boldsymbol{x}^\top \boldsymbol{e}_{\mathscr{T}}$ only takes the element in $\boldsymbol{x}$ that corresponds to $\mathscr{T}$. Suppose $\mathscr{T}_{m'} = \mathscr{T}$ for some $0 \le m' \le L$. By Supplementary Eq. (11), $x_{m'}$ is independent of the entries in $V$ that are associated with trees $\mathscr{T}_l$ for $l > m'$. Moreover, by the definition of $Q$ in Supplementary Eq. (2), $Q_{\mathscr{T}_l,\mathscr{T}_{m'}}$ is non-zero only if $\mathscr{T}_l$ is a subtree of $\mathscr{T}_{m'}$, *i.e.* $\mathscr{T}_l \subset \mathscr{T}_{m'}$. Therefore, in order to compute $x_{m'}$, we only need a sub-matrix $Q_{\mathscr{T}}$ of $Q$, where the columns and rows correspond to all subtrees of $\mathscr{T}$ ordered by increasing size (Supplementary Figure S1(c)),

$$Q_{\mathscr{T}} = (q_{\mathscr{T}_1,\mathscr{T}_2} : \mathscr{T}_1, \mathscr{T}_2 \subseteq \mathscr{T}). \tag{12}$$

In particular, the first and the last row (column) of $Q_{\mathscr{T}}$ correspond to $\mathscr{T}_0$ and $\mathscr{T}$ respectively. Hence, Supplementary Eq. (6) becomes

$$p(\mathscr{T} \mid \Theta) = \lambda_s (\underbrace{\lambda_s I - Q_{\mathscr{T}}}_{V_{\mathscr{T}}})^{-1}_{\mathscr{T}_0,\mathscr{T}}, \tag{13}$$

where the dimension of $V_{\mathscr{T}}$ is determined by the number of subtrees of $\mathscr{T}$, which is much less than $L$. Note that since $Q_{\mathscr{T}}$ is often very sparse, a sparse triangular matrix solver can be helpful.

Next, the gradient with respect to each $\Theta_{ij}$ can be computed as

$$\begin{aligned}
\frac{\partial \log p(\mathscr{T} \mid \Theta)}{\partial \Theta_{ij}} &= \frac{1}{p(\mathscr{T} \mid \Theta)} \frac{\partial p(\mathscr{T} \mid \Theta)}{\partial \Theta_{ij}} && \text{by (PP136)} \\
&= \frac{1}{p(\mathscr{T} \mid \Theta)} \frac{\partial}{\partial \Theta_{ij}} \left[ \lambda_s (V_{\mathscr{T}})^{-1}_{\mathscr{T}_0,\mathscr{T}} \right] && \text{by Supplementary Eq. (13)} \\
&= \frac{\lambda_s}{p(\mathscr{T} \mid \Theta)} \left[ \frac{\partial V_{\mathscr{T}}^{-1}}{\partial \Theta_{ij}} \right]_{\mathscr{T}_0,\mathscr{T}} && \text{by (PP32) and (PP34)} \\
&= \frac{\lambda_s}{p(\mathscr{T} \mid \Theta)} \left[ -V_{\mathscr{T}}^{-1} \frac{\partial V_{\mathscr{T}}}{\partial \Theta_{ij}} V_{\mathscr{T}}^{-1} \right]_{\mathscr{T}_0,\mathscr{T}} && \text{by (PP59)} \\
&= \frac{\lambda_s}{p(\mathscr{T} \mid \Theta)} \left[ -V_{\mathscr{T}}^{-1} \Big( \sum_{(\pi,i): j \in \pi \text{ or } j=i} \frac{\partial V_{\mathscr{T}}}{\partial \lambda_{(\pi,i)}} \frac{\partial \lambda_{(\pi,i)}}{\partial \Theta_{ij}} \Big) V_{\mathscr{T}}^{-1} \right]_{\mathscr{T}_0,\mathscr{T}} && \text{by (PP35) and (PP136)} \\
&= \frac{\lambda_s}{p(\mathscr{T} \mid \Theta)} \left[ \sum_{(\pi,i): j \in \pi \text{ or } j=i} -V_{\mathscr{T}}^{-1} \frac{\partial V_{\mathscr{T}}}{\partial \lambda_{(\pi,i)}} \frac{\partial \lambda_{(\pi,i)}}{\partial \Theta_{ij}} V_{\mathscr{T}}^{-1} \right]_{\mathscr{T}_0,\mathscr{T}} && \text{by the distributive law} \\
&= \frac{1}{p(\mathscr{T} \mid \Theta)} \sum_{(\pi,i): j \in \pi \text{ or } j=i} \underbrace{\lambda_s \Big( -V_{\mathscr{T}}^{-1} \frac{\partial V_{\mathscr{T}}}{\partial \lambda_{(\pi,i)}} V_{\mathscr{T}}^{-1} \Big)_{\mathscr{T}_0,\mathscr{T}}}_{\partial p(\mathscr{T} \mid \Theta) / \partial \lambda_{(\pi,i)}} \frac{\partial \lambda_{(\pi,i)}}{\partial \Theta_{ij}} && \text{by (PP32) and (PP34)}
\end{aligned}$$

where the numbered properties can be found in the Matrix Cookbook by Petersen and Pedersen, 2012*. We add "PP" in front the numbers to distinguish between the numbers in the Matrix Cookbook and the

*https://www.math.uwaterloo.ca/~hwolkowi/matrixcookbook.pdf

numbers of the equations in this paper. The term $\partial p(\mathscr{T} \mid \Theta)/\partial\lambda_{(\pi,i)}$ can be obtained from the first row and the last column of the matrix

$$\lambda_s(\lambda_s I - R_{\mathscr{T}}^{(\pi,i)})^{-1}, \qquad R_{\mathscr{T}}^{(\pi,i)} := \begin{bmatrix} Q_{\mathscr{T}} & \frac{\partial Q_{\mathscr{T}}}{\partial \lambda_{(\pi,i)}} \\ 0 & Q_{\mathscr{T}} \end{bmatrix},$$

with forward substitution, because

$$(\lambda_s I - R_{\mathscr{T}}^{(\pi,i)})^{-1} = \begin{bmatrix} \lambda_s I - Q_{\mathscr{T}} & -\frac{\partial Q_{\mathscr{T}}}{\partial \lambda_{(\pi,i)}} \\ 0 & \lambda_s I - Q_{\mathscr{T}} \end{bmatrix}^{-1} = \begin{bmatrix} V_{\mathscr{T}} & \frac{\partial V_{\mathscr{T}}}{\partial \lambda_{(\pi,i)}} \\ 0 & V_{\mathscr{T}} \end{bmatrix}^{-1} = \begin{bmatrix} V_{\mathscr{T}}^{-1} & -V_{\mathscr{T}}^{-1}\frac{\partial V_{\mathscr{T}}}{\partial \lambda_{(\pi,i)}} V_{\mathscr{T}}^{-1} \\ 0 & V_{\mathscr{T}}^{-1} \end{bmatrix}$$

by (PP399) and (PP400) of the Matrix Cookbook. Also, the gradient of the penalty term $\gamma \sum_{i \neq j} |\log \Theta_{ij}|$ with respect to each $\Theta_{ij}$ is

$$\frac{\gamma \operatorname{sign}(\theta_{ij})}{\Theta_{ij}}, \qquad \operatorname{sign}(\theta_{ij}) = \begin{cases} 1 & \theta_{ij} > 0 \\ 0 & \theta_{ij} = 0 \\ -1 & \theta_{ij} < 0 \end{cases}.$$

Therefore, given $N$ mutation trees $\boldsymbol{\mathscr{T}} = \{\mathscr{T}^1, \ldots, \mathscr{T}^N\}$, we can find the maximum likelihood estimation of $\Theta$ by gradient ascent optimization.

#### A.4 Hybrid EM and Monte Carlo EM algorithm for the estimation of an MHN given a set of trees

The joint density of a collection of waiting times $\boldsymbol{t}$, sampling time $t_s$, and the corresponding tree structure $\mathscr{T}$ is

$$\begin{aligned} p(\mathscr{T}, \boldsymbol{t}, t_s \mid \Theta) &= f_{\lambda_s}(t_s) \prod_{i=1}^{n} \prod_{\pi \in \mathscr{T}:(\pi,i) \in A(\mathscr{T})} f_{\lambda_{(\pi,i)}}(t_{(\pi,i)} - t_\pi) \times \prod_{\pi \in \mathscr{T}} \mathbb{1}(t_\pi < t_s) \times \prod_{\pi \in A(\mathscr{T}):\pi \notin \mathscr{T}} \mathbb{1}(t_\pi > t_s) \\ &= f_{\lambda_s}(t_s) \prod_{i=1}^{n} \prod_{\pi \in \mathscr{T}:(\pi,i) \in A(\mathscr{T})} \Big[\lambda_{(\pi,i)} \exp\big(-\lambda_{(\pi,i)}(t_{(\pi,i)} - t_\pi)\big)\Big] \\ &\quad \times \prod_{\pi \in \mathscr{T}} \mathbb{1}(t_\pi < t_s) \times \prod_{\pi \in A(\mathscr{T}):\pi \notin \mathscr{T}} \mathbb{1}(t_\pi > t_s) \\ &= f_{\lambda_s}(t_s) \prod_{i=1}^{n} \prod_{\pi \in \mathscr{T}:(\pi,i) \in A(\mathscr{T})} \Big[\Theta_{ii} \prod_{j \in \pi} \Theta_{ij} \exp\big(-\Theta_{ii} \prod_{j \in \pi} \Theta_{ij}(t_{(\pi,i)} - t_\pi)\big)\Big] \\ &\quad \times \prod_{\pi \in \mathscr{T}} \mathbb{1}(t_\pi < t_s) \times \prod_{\pi \in A(\mathscr{T}):\pi \notin \mathscr{T}} \mathbb{1}(t_\pi > t_s) \end{aligned}$$

where $f_\lambda$ is the exponential density function with parameter $\lambda$ and $\lambda_s = 1$. The complete-data log-likelihood is then

$$\ell^{\text{full}}(\Theta) = \sum_{l=1}^{N} \sum_{i=1}^{n} \sum_{\pi \in \mathscr{T}^l:(\pi,i) \in A(\mathscr{T}^l)} \Big[\log \Theta_{ii} + \sum_{j \in \pi} \log \Theta_{ij} - \Theta_{ii} \prod_{j \in \pi} \Theta_{ij}(t^l_{(\pi,i)} - t^l_\pi)\Big] + \log f_{\lambda_s}(t^l_s)$$

and $\log f_{\lambda_s}(t^l_s)$ is a constant with respect to $\Theta$. The expected complete-data log-likelihood in the E step of the EM algorithm follows by replacing each $(t_{(\pi,i)} - t_\pi)$ with $\mathbb{E}_{\boldsymbol{T}, T_s \mid \mathscr{T}, \Theta}[T_{(\pi,i)} - T_\pi]$.

To compute the expected time differences, we first use the definition of the expected value,

$$\begin{aligned} \mathbb{E}_{\boldsymbol{T}, T_s \mid \mathscr{T}, \Theta}[T_{(\pi,i)} - T_\pi] &= \int_{\mathbb{R}_{\geq 0}^{|A(\mathscr{T})|+1}} (t_{(\pi,i)} - t_\pi) p(\boldsymbol{t}, t_s \mid \mathscr{T}, \Theta) d\boldsymbol{t} dt_s \\ &= \frac{1}{P(\mathscr{T} \mid \Theta)} \int_{\mathbb{R}_{\geq 0}^{|A(\mathscr{T})|+1}} (t_{(\pi,i)} - t_\pi) p(\mathscr{T}, \boldsymbol{t}, t_s \mid \Theta) d\boldsymbol{t} dt_s. \end{aligned}$$

Notice that

$$\begin{aligned}\frac{\partial p(\mathscr{T} \mid \Theta)}{\partial \lambda_{(\pi,i)}} &= \frac{\partial}{\partial \lambda_{(\pi,i)}} \int_{\mathbb{R}_{\geq 0}^{|A(\mathscr{T})|+1}} p(\mathscr{T}, \boldsymbol{t}, t_s \mid \Theta) d\boldsymbol{t} dt_s \\ &= \int_{\mathbb{R}_{\geq 0}^{|A(\mathscr{T})|+1}} \frac{\partial}{\partial \lambda_{(\pi,i)}} p(\mathscr{T}, \boldsymbol{t}, t_s \mid \Theta) d\boldsymbol{t} dt_s \\ &= \int_{\mathbb{R}_{\geq 0}^{|A(\mathscr{T})|+1}} \Big(\frac{1}{\lambda_{(\pi,i)}} - (t_{(\pi,i)} - t_\pi)\Big) p(\mathscr{T}, \boldsymbol{t}, t_s \mid \Theta) d\boldsymbol{t} dt_s \\ &= p(\mathscr{T} \mid \Theta)\Big(\frac{1}{\lambda_{(\pi,i)}} - \mathbb{E}_{\boldsymbol{T}, T_s \mid \mathscr{T}, \Theta}[T_{(\pi,i)} - T_\pi]\Big).\end{aligned}$$

Therefore,

$$\mathbb{E}_{\boldsymbol{T}, T_s \mid \mathscr{T}, \Theta}[T_{(\pi,i)} - T_\pi] = \frac{1}{\lambda_{(\pi,i)}} - \frac{1}{P(\mathscr{T} \mid \Theta)} \frac{\partial P(\mathscr{T} \mid \Theta)}{\partial \lambda_{(\pi,i)}},$$

where both $P(\mathscr{T} \mid \Theta)$ and $\frac{\partial P(\mathscr{T} \mid \Theta)}{\partial \lambda_{(\pi,i)}}$ can be computed as in Section A.3. For large trees (*e.g.* with more than 500 subtrees), however, computing this exact form can be very slow. In this case, we can approximate it by drawing $M$ samples from the proposal distribution with density $g$ as defined in Methods. To recapitulate, we first draw the sampling time $T_s$ from $\mathrm{Exp}(\lambda_s)$ with $\lambda_s = 1$, followed by drawing the difference in waiting times between subclones $\pi$ and $(\pi, i)$ recursively as

$$Z_{(\pi,i)} \sim \begin{cases} \mathrm{TExp}(\lambda_{(\pi,i)}, 0, t_s - t_\pi) & \text{if } (\pi, i) \in \mathscr{T} \\ \mathrm{TExp}(\lambda_{(\pi,i)}, t_s - t_\pi, \infty) & \text{if } (\pi, i) \in A(\mathscr{T}) \setminus \mathscr{T} \end{cases}$$

where $\mathrm{TExp}(\lambda, a, b)$ is the truncated exponential distribution with parameter $\lambda$ and bounds $0 \leq a < b < \infty$. The importance sampling weights are defined as

$$w^{(m)} := w(\boldsymbol{t}^{(m)} \mid \mathscr{T}, \Theta) = \frac{p(\mathscr{T}, \boldsymbol{t}^{(m)}, t_s^{(m)} \mid \Theta)}{g(\boldsymbol{t}^{(m)}, t_s^{(m)} \mid \mathscr{T}, \Theta)}, \qquad m = 1, \ldots, M.$$

The densities of the truncated exponential distributions $\mathrm{TExp}(\lambda, 0, a)$ and $\mathrm{TExp}(\lambda, b, \infty)$ are

$$f_{\lambda,0,a}^{\mathrm{TExp}}(t) = \frac{\lambda e^{-\lambda t} \mathbb{1}_{[0,a]}(t)}{1 - e^{-\lambda a}} \qquad \text{and} \qquad f_{\lambda,b,\infty}^{\mathrm{TExp}}(t) = \frac{\lambda e^{-\lambda t} \mathbb{1}_{[b,\infty)}(t)}{e^{-\lambda b}} \qquad \text{respectively.}$$

Since all terms in the numerator of $w^{(m)}$ cancel out, it follows that

$$w^{(m)} = \prod_{i=1}^{n} \prod_{(\pi,i) \in \mathscr{T}} (1 - e^{-\lambda_{(\pi,i)}(t_s^{(m)} - t_\pi^{(m)})}) \prod_{(\pi,i) \in A(\mathscr{T}),\ (\pi,i) \notin \mathscr{T}} e^{-\lambda_{(\pi,i)}(t_s^{(m)} - t_\pi^{(m)})}.$$

The approximation becomes

$$\mathbb{E}_{\boldsymbol{T}, T_s \mid \mathscr{T}, \Theta}[T_{(\pi,i)} - T_\pi] \approx \frac{\frac{1}{M} \sum_{m=1}^{M} w^{(m)} (t_{(\pi,i)}^{(m)} - t_\pi^{(m)})}{\frac{1}{M} \sum_{m=1}^{M} w^{(m)}}.$$

In the M step, we update $\Theta$ by maximizing the penalized expected complete-data log-likelihood, where the gradients with respect to each $\Theta_{ij}$ are computed as follows:

$$\begin{cases} \frac{\partial Q(\Theta, \Theta^{(k)})}{\partial \Theta_{ii}} = \sum_{l=1}^{N} \sum_{\pi \in \mathscr{T}^l : (\pi,i) \in A(\mathscr{T}^l)} \Big[\frac{1}{\Theta_{ii}} - \prod_{j \in \pi} \Theta_{ij} \mathbb{E}_{\boldsymbol{T}^l, T_s^l \mid \mathscr{T}^l, \Theta^{(k)}}[T_{(\pi,i)}^l - T_\pi^l]\Big] & i \in [n] \\ \frac{\partial Q(\Theta, \Theta^{(k)})}{\partial \Theta_{ij}} = \sum_{l=1}^{N} \sum_{\pi \in \mathscr{T}^l : (\pi,i) \in A(\mathscr{T}^l), j \in \pi} \Big[\frac{1}{\Theta_{ij}} - \Theta_{ii} \prod_{k \in \pi : k \neq j} \Theta_{ik} \mathbb{E}_{\boldsymbol{T}^l, T_s^l \mid \mathscr{T}^l, \Theta^{(k)}}[T_{(\pi,i)}^l - T_\pi^l]\Big] & i, j \in [n],\ i \neq j \end{cases}.$$

#### A.5 Parameter estimation based on non-empty trees

In practice, we typically cannot observe an empty mutation tree containing only the root node for a cancer patient and all observed trees in $\boldsymbol{\mathscr{T}}$ have at least one mutation. In this case, we need to condition the likelihood on the trees being non-empty, which amounts to adding the following term to the objective functions in both the MLE and the EM parameter estimation procedure:

$$-N \log(1 - p(\mathscr{T}_0 \mid \Theta)) = -N\Big(1 - \frac{\lambda_s}{\lambda_s + \sum_{i=1}^n \Theta_{ii}}\Big).$$

To see this, consider the marginal probability of a tree $\mathscr{T}$ given $\Theta$. It can be decomposed into two parts,

$$\begin{aligned} p(\mathscr{T} \mid \Theta) &= p(\mathscr{T}, \mathscr{T} \text{ is empty} \mid \Theta) + p(\mathscr{T}, \mathscr{T} \text{ is non-empty} \mid \Theta) \\ &= p(\mathscr{T} \mid \mathscr{T} \text{ is empty}, \Theta) p(\mathscr{T} \text{ is empty} \mid \Theta) \mathbb{1}(\mathscr{T} \text{ is empty}) \\ &\quad + p(\mathscr{T} \mid \mathscr{T} \text{ is non-empty}, \Theta) p(\mathscr{T} \text{ is non-empty} \mid \Theta) \mathbb{1}(\mathscr{T} \text{ is non-empty}). \end{aligned}$$

Since $\mathscr{T}$ can be either empty or non-empty, only one term in the summation is left. It follows that

$$p(\mathscr{T} \mid \mathscr{T} \text{ is non-empty}, \Theta) = \frac{p(\mathscr{T} \mid \Theta)}{p(\mathscr{T} \text{ is non-empty} \mid \Theta)} = \frac{p(\mathscr{T} \mid \Theta)}{1 - p(\mathscr{T} \text{ is empty} \mid \Theta)}.$$

Now, $\mathscr{T}$ is empty if and only if $T_s = \min\{T_s, T_{(0,1)}, \ldots, T_{(0,n)}\}$, which is the minimum of a set of independent exponential random variables. Therefore,

$$p(\mathscr{T} \text{ is empty} \mid \Theta) = \frac{\lambda_s}{\lambda_s + \lambda_{(0,1)} + \cdots + \lambda_{(0,n)}} = \frac{\lambda_s}{\lambda_s + \sum_{i=1}^n \Theta_{ii}},$$

and

$$\frac{\partial}{\partial \Theta_{ii}} \log(1 - p(\mathscr{T} \text{ is empty} \mid \Theta)) = \frac{p(\mathscr{T} \text{ is empty} \mid \Theta)^2}{\lambda_s (1 - p(\mathscr{T} \text{ is empty} \mid \Theta))}.$$

Finally, the MLE objective function becomes

$$\begin{aligned} \hat{\Theta} &= \arg\max_{\Theta} \sum_{l=1}^N \log p(\mathscr{T}^l \mid \mathscr{T} \text{ is non-empty}, \Theta) - \gamma \sum_{i \neq j} |\log \Theta_{ij}| \\ &= \arg\max_{\Theta} \sum_{l=1}^N \log p(\mathscr{T}^l \mid \Theta) - \gamma \sum_{i \neq j} |\log \Theta_{ij}| - N\Big(1 - \frac{\lambda_s}{\lambda_s + \sum_{i=1}^n \Theta_{ii}}\Big). \end{aligned}$$

Note that the same term can also be added to the objective function of the genotype MHN method if none of the input genotypes correspond to the wild type (*i.e.* a vector of zeros).

#### B TreeMHN with stability selection

TreeMHN can be viewed as a probabilistic graphical model with $n^2$ parameters, where the $(n^2 - n)$ off-diagonal entries represent the edges ($\rightarrow$ or $\dashv$) among the mutations (Figure S1 (a)). When the sample size $N$ is small, it is often difficult to identify the true edges. In this case, we use a procedure called stability selection, which is particularly suitable for controlling the false discovery rate in high-dimensional graphical modelling problems [3].

More specifically, we take a random subsample $I$ of size $\lfloor \frac{N}{2} \rfloor$ without replacement from $\mathscr{T}$ and train TreeMHN for a given regularization parameter $\gamma > 0$. An off-diagonal entry $\theta_{ij}$ is considered as selected if it is non-zero. The set of selected edges is

$$\hat{S}_\gamma(I) = \{(i,j) \mid |\theta_{ij}| > 0, i \neq j\}. \tag{14}$$

Repeating this process for $B$ times ($B$ large), the probability that an entry at position $(i,j)$ for $i \neq j$ is selected can be approximated as

$$\hat{q}_{(i,j)}(\gamma) = \frac{1}{B} \sum_{b=1}^{B} \mathbb{1}\{(i,j) \in \hat{S}_\gamma(I^b)\}. \tag{15}$$

For a pre-specified threshold $\delta$ and a regularization parameter $\gamma$, the set of stable edges is defined as [3]

$$\hat{S}_{\text{stable}}(\gamma) = \{(i,j) \mid \hat{q}_{(i,j)}(\gamma) \geq \delta\}. \tag{16}$$

Let $S_0$ be the set of true edges, $S_0^c$ be the complement of $S_0$, and $V = |S_0^c \cap \hat{S}_{\text{stable}}|$ be the falsely selected edges. To control for the expected number of false positives (*e.g.* $\mathbb{E}[V] \leq \nu = 5$), we choose to fix the threshold $\delta$ (*e.g.* 0.95) and vary $\gamma$ such that the number of stable edges is bounded (Meinshausen and Bühlmann, 2010),

$$|\hat{S}_{\text{stable}}(\gamma)| \leq \lfloor \sqrt{\nu(n^2 - n)(2\delta - 1)} \rfloor. \tag{17}$$

After estimating $\hat{S}_{\text{stable}}(\gamma)$, we can then refit TreeMHN on the full dataset $\mathscr{T}$ by masking the entries in $\Theta$ that are not selected.

### C Computing probabilities of trajectories and mutational events with TreeMHN

#### C.1 Most probable evolutionary trajectories

In Methods, we have introduced how to compute the trajectory probability distribution over $\Pi_S$, the set of evolutionary trajectories that end with the sampling event, and $\Pi_d$, the set of evolutionary trajectories of a fixed length $d$. There are advantages and limitations associated with both formulations. Using trajectory probabilities over $\Pi_d$, we can compare the inferred trajectory distribution from TreeMHN with the ones from methods that do not have the notion of a sampling event. It is also easy to compute for small $d$. However, we cannot compare trajectories of different lengths or with the frequencies of the trajectories observed in the tumor trees based on this formulation. The trajectory probability distribution over $\Pi_S$, on the other hand, does not have these problems, but it can become infeasible to compute for even a moderate number of mutations $n$, because the space grows super-exponentially with $n$. Instead, we can enumerate the most probable trajectories that have at least one mutation dynamically with the pseudocode in Algorithm 2.

---

**Algorithm 2** Most probable evolutionary trajectories

---

**Input**

| | |
|---|---|
| $\Theta$ | Mutual Hazard Network |
| $\tau$ | Number of most probable trajectories to output |
| $\lambda_s$ | Sampling rate |

**Output**

| | |
|---|---|
| $\Pi_\tau := \{\pi_{(1)}, \ldots, \pi_{(\tau)}\}$ | The top $\tau$ most probable trajectories |
| $\mathbf{p}_\tau := \{p_{(1)}, \ldots, p_{(\tau)}\}$ | Probabilities of the $\tau$ most probable trajectories |

```
Π_τ ← {(0), ..., (0)}                      ▷ Initialize to trajectories with no mutations
p_τ ← (0, ..., 0)                          ▷ Initialize to zero for all trajectory probabilities
Π_current ← {(0,1), ..., (0,n)}            ▷ trajectories to visit
while Π_current is non-empty do
    Π_next ← {}                            ▷ Initialize the set of trajectories to visit next
    for π in Π_current do
        p_π ← probability of π computed using Eq. (10) in the main text
        if p_π > min(p_τ) then
            r ← index of the first minimum in p_τ
            π_(r) ← π                      ▷ Replace the r^th most probable trajectory by π
            p_(r) ← p_π
            Π_ch(π) ← {(π, j) | j ∈ [n] \ π}   ▷ Children trajectories of π by extending π by one mutation
            Π_next ← {Π_next, Π_ch(π)}     ▷ Append the children trajectories to the trajectories to visit next
        end if
        Π_current ← Π_next
    end for
end while
p_τ ← p_τ/(1 − λ_s/(λ_s + Σ_{j∈[n]} λ_(0,j)))   ▷ Conditioned on trajectories that have at least one mutation
```

---

#### C.2 Alternative methods to compute the probabilities of mutational events

Although TreeMHN is unique in its ability to compute the probabilities of the next mutational events given a tree (Methods), we adapt five alternative methods for comparison:

1. **A frequency-based model**: A frequency-based model ($N_0$) predicts the next event using the relative frequencies of the mutations in the cohort, $(f_1, \ldots, f_n)$ with $f_i > 0$ and $\sum_{i=1}^{n} f_i = 1$. The new event can be placed randomly after any node (including the root) in $\mathscr{T}$ with probability $1/|\mathscr{T}|$. Then, the

conditional probability of an event $\pi = (0, \sigma_1, \ldots, \sigma_d) \in A(\mathscr{T}) \setminus \mathscr{T}$ happening before all the other events depends only on the relative frequency of $\sigma_d$ and is given by

$$p(\pi \mid \mathscr{T}, N_0) = \frac{1}{|\mathscr{T}|} \times f_{\sigma_d} \times \mathbb{1}\{\pi \in A(\mathscr{T}) \setminus \mathscr{T}\} \times \frac{1}{Z(\mathscr{T})}, \quad \sigma_d \in [n], \quad 1 \leq d \leq n, \tag{18}$$

where $Z(\mathscr{T})$ is a normalizing constant depending on $\mathscr{T}$ such that $\sum_{\pi \in A(\mathscr{T}) \setminus \mathscr{T}} p(\pi \mid \mathscr{T}, N_0) = 1$. We can estimate the relative frequencies $(\hat{f}_1, \ldots, \hat{f}_n)$ using the subclonal genotypes weighted by their relative sizes (Supplementary Figure S2).

2. **TreeMHN with only the baseline rates**: We can run TreeMHN on a set of mutation trees with the restriction that all off-diagonal elements in the estimated network are zero. In other words, this approach assumes that all mutations are independent of each other while respecting the tree structures. The ways to compute the trajectory probabilities and the probabilities of the next mutational events given a tree stay the same.

3. **Genotype MHN using the consensus genotypes**: We can run the genotype MHN method of [4] on the consensus genotypes of patient samples (Supplementary Figure S2). Then, we use the estimated network to compute the trajectory probabilities and the probabilities of the next mutational events under the TreeMHN framework.

4. **Genotype MHN using the weighted subclonal genotypes**: Same as 2 except that we use the subclonal genotypes weighted by the size of the subclones (Supplementary Figure S2).

5. **REVOLVER**: The row-normalized matrix $\boldsymbol{w}$ in REVOLVER summarizes the relative frequency of mutation $j$ being the descendant of $i$ in its entry $w_{ij}$. Given $\boldsymbol{w}$, the probability of a new event with mutation $j$ given a tree $\mathscr{T}$ can be computed as

$$\frac{1}{|\mathscr{T}|} \times w_{ij} \times \mathbb{1}\{j \text{ is placed after a node } i\} \times \frac{1}{Z(\mathscr{T})}, \tag{19}$$

where $Z(\mathscr{T})$ is a normalizing constant depending on $\mathscr{T}$ such that the probabilities sum up to 1 over all possible events.

| Sample | Subclone size | Mutation 1 | Mutation 2 | Mutation 3 |
|---|---|---|---|---|
| A | **0.55** | 1 | 0 | 0 |
| | **0.3** | 0 | 1 | 0 |
| | **0.15** | 0 | 1 | 1 |
| B | **0.38** | 1 | 0 | 1 |
| | **0.62** | 1 | 1 | 0 |
| C | **0.2** | 1 | 0 | 0 |
| | **0.35** | 0 | 1 | 1 |
| | **0.18** | 1 | 1 | 0 |
| | **0.27** | 1 | 0 | 0 |

(a)

| Sample | Consensus (> 50%) | | |
|---|---|---|---|
| **A** | 1 | 0 | 0 |
| **B** | 1 | 1 | 0 |
| **C** | 1 | 1 | 0 |

(b)

| Mutation 1 | Mutation 2 | Mutation 3 |
|---|---|---|
| 0.47 | 0.34 | 0.19 |

(c)

Figure S2: An example of converting subclonal genotypes (a) to consensus genotypes (b) using a threshold of 50%. The relative frequencies of the three mutations are shown in (c).

### D Additional simulation details

#### D.1 Precision and recall of identifying the true MHN

In the objective functions of both TreeMHN and the genotype MHN method, there is a penalization parameter $\gamma$ that controls the sparsity of the estimated network $\hat{\Theta}$. The larger $\gamma$ is, the more zero entries are in $\hat{\Theta}$. We evaluate the structural differences between $\hat{\Theta}$ and a ground truth network $\Theta$ by defining the true positives (TP), false positives (FP), true negatives (TN), and false negatives (FN) as indicated in Figure S3. In particular, an edge in $\hat{\Theta}$ is considered as a TP if and only if it has the correct direction ($\text{sign}(\hat{\theta}_{ij}) = \text{sign}(\theta_{ij})$). Next, we define the precision and recall as

$$\text{Precision} = \frac{\text{TP}}{\hat{\text{P}}} \quad \text{and} \quad \text{Recall} = \frac{\text{TP}}{\text{P}}, \tag{20}$$

where P and $\hat{\text{P}}$ are the number of edges in the true network $\Theta$ and the estimated network $\hat{\Theta}$ respectively. The $F_1$ score is the harmonic mean between precision and recall, which can be computed as

$$\text{F1 score} = 2 \times \frac{\text{Precision} \times \text{Recall}}{\text{Precision} + \text{Recall}}. \tag{21}$$

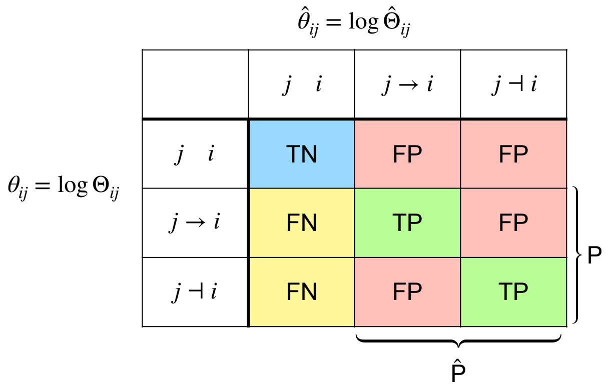

Figure S3: Definition of true positives (TP), false positives (FP), true negatives (TN), and false negatives (FN) for the structural differences between an estimated network $\hat{\Theta}$ and a true network $\Theta$.

On average, a network estimated by randomly guessing the direction of the edges can only achieve a maximum recall of 50%. The precision of random guess is equal to $0.5(1 - s)$, where $s$ is the proportion of zero off-diagonal entries in $\Theta$, also called network sparsity. If 50% of the entries in $\Theta$ are zero, then the precision is only 25%. The corresponding $F_1$ score is $\frac{1}{3}$.

#### D.2 Computing trajectory probabilities with REVOLVER and HINTRA

REVOLVER [5] and HINTRA [6] are two probabilistic methods for inferring repeated trajectories from cross-sectional multi-region sequencing data. Both methods learn one tree for each patient by transferring the information across tumors, which is summarized in matrix $\boldsymbol{w}$ and $\beta$, respectively. For REVOLVER, $\boldsymbol{w}$ is an $n$-by-$n$ matrix, where an entry $w_{ij}$ is the number of times mutation $i$ occurs before mutation $j$ in the trees. By row-normalizing $\boldsymbol{w}$, one can obtain the empirical probability of mutation $j$ being the descendant of mutation $i$. For HINTRA, the columns of $\beta$ also correspond to the $n$ mutations, but the rows are all possible ancestry sets, which increases super-exponentially in $n$. To obtain the matrices $\boldsymbol{w}$ and $\beta$, we can count the frequency of every possible ancestor-descendant pair that appears in the simulated trees. Since we only consider evolutionary trajectories of length 4, it is still feasible to compute the HINTRA matrix $\beta$. Adding pseudocounts and row-normalizing the two matrices gives us $\tilde{\boldsymbol{w}}$ and $\tilde{\beta}$, which can be used to estimate trajectory probabilities. Given $\tilde{\boldsymbol{w}}$ or $\tilde{\beta}$, one simply multiplies the entries in the matrices that correspond to the edges in the trajectories. For example, if an evolutionary trajectory is $A \to B \to C$, then we have $\tilde{w}_{\emptyset,A} \times \tilde{w}_{A,B} \times \tilde{w}_{B,C}$ for REVOLVER and $\tilde{\beta}_{\emptyset,A} \times \tilde{\beta}_{\{A\},B} \times \tilde{\beta}_{\{A,B\},C}$ for HINTRA. Moreover, REVOLVER requires an additional step of normalizing over all trajectories in $\Pi_d$ to ensure that the probabilities sum up to 1.

#### D.3 Introducing noise into simulated trees

To evaluate the robustness of TreeMHN against tree uncertainty, we define a noise level $\epsilon \in (0, 1)$ and use it to perturb the tree structures. For each node except the root in a tree, we generate a random number $r \sim \text{Uniform}(0, 1)$ and perturb the node if $r < \epsilon$. Depending on which of the following categories the node falls into, we randomly choose one way of perturbation:

- Internal node with multiple children:
  - randomly insert a parent node (the original parent is shifted up);
  - randomly add a child node.
- Internal node with exactly one child:
  - randomly insert a parent node;
  - randomly add a child node;
  - switch order with the child;
  - be removed.
- Leaf node:
  - randomly insert a parent node;
  - randomly add a child node;
  - prune the node and reattach it to one of its siblings or the parent of its parent;
  - be removed.

To remove a node, we also need to ensure that it is not the only node in the tree.

#### D.4 Additional figures from simulations

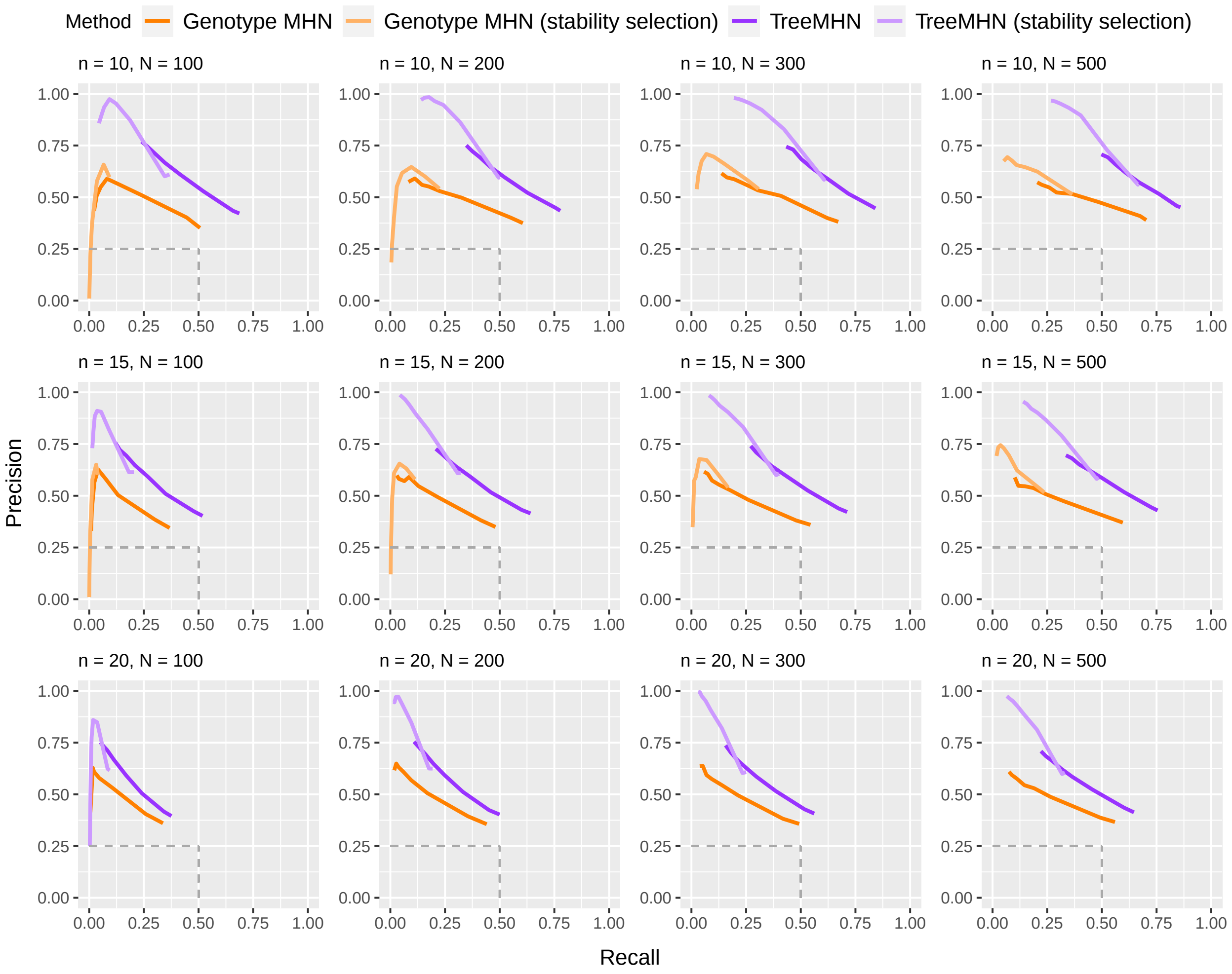

Figure S4: Performance of TreeMHN and the genotype MHN methods (with and without stability selection) in estimating the true network $\Theta$ on simulated data for $n \in \{10, 15, 20\}$ and $N \in \{200, 300, 500\}$. The precision and recall curves averaged over 100 simulation runs are plotted by comparing the full matrices $\Theta$ and $\hat{\Theta}$ (see Figure 3 for comparing sub-matrices restricted to top 50% of the mutations ranked by baseline rates). Each point on the curves corresponds to a penalization level $\gamma \in \{0.05, 0.1, 0.5, 1, 1.5, 2, 2.5, 3\}$. The dash lines indicate the performance of randomly guessing the edge directions in $\Theta$.

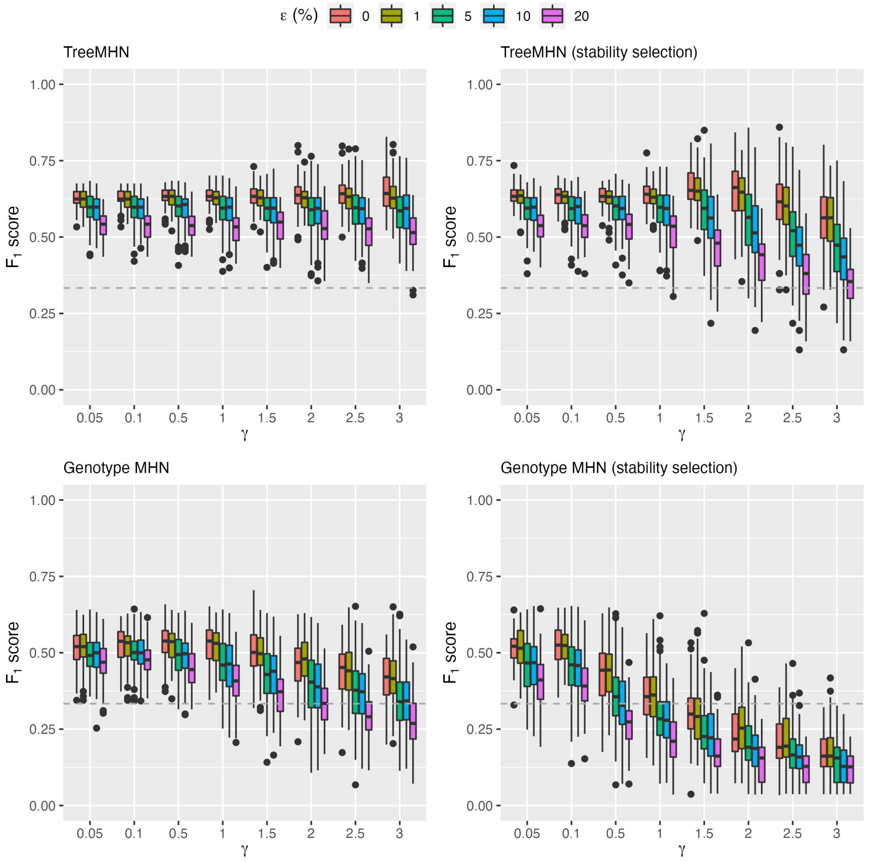

Figure S5: Performance of TreeMHN and the genotype MHN method (with and without stability selection) on perturbed simulated trees at noise level $\epsilon \in \{1\%, 5\%, 10\%, 20\%\}$ for $n = 10$ and $N = 500$. For each method, we plot the $F_1$ score over 100 simulation runs at $\gamma \in \{0.05, 0.1, 0.5, 1, 1.5, 2, 2.5, 3\}$. $F_1$ score measures the harmonic mean between precision and recall. In general, $F_1$ score decreases as noise level increases.

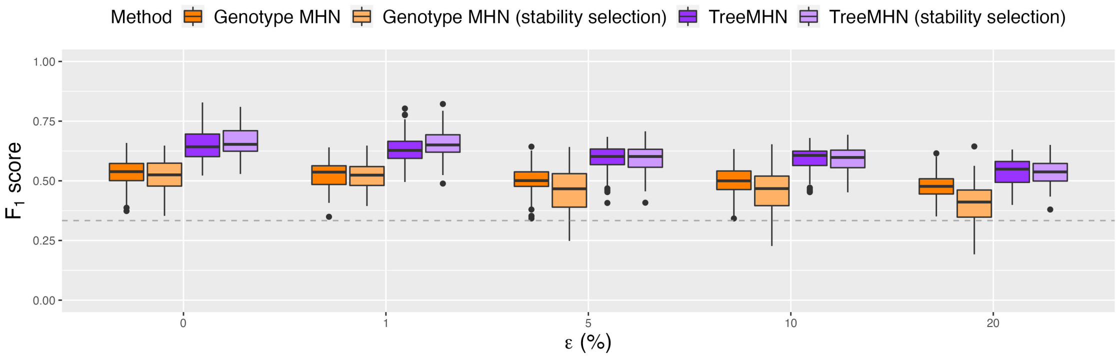

Figure S6: Performance of TreeMHN and the genotype MHN method (with and without stability selection) in estimating the true network $\Theta$ on perturbed simulated trees at noise level $\epsilon \in \{0\%, 1\%, 5\%, 10\%, 20\%\}$ for $n = 10$ and $N = 500$. $F_1$ score measures the harmonic mean between precision and recall. For each method and each $\epsilon$, the box contains 100 simulation runs and corresponds to the optimal regularization parameter with respect to the average $F_1$ score (Figure S5). The dash line indicates the performance of random guess.

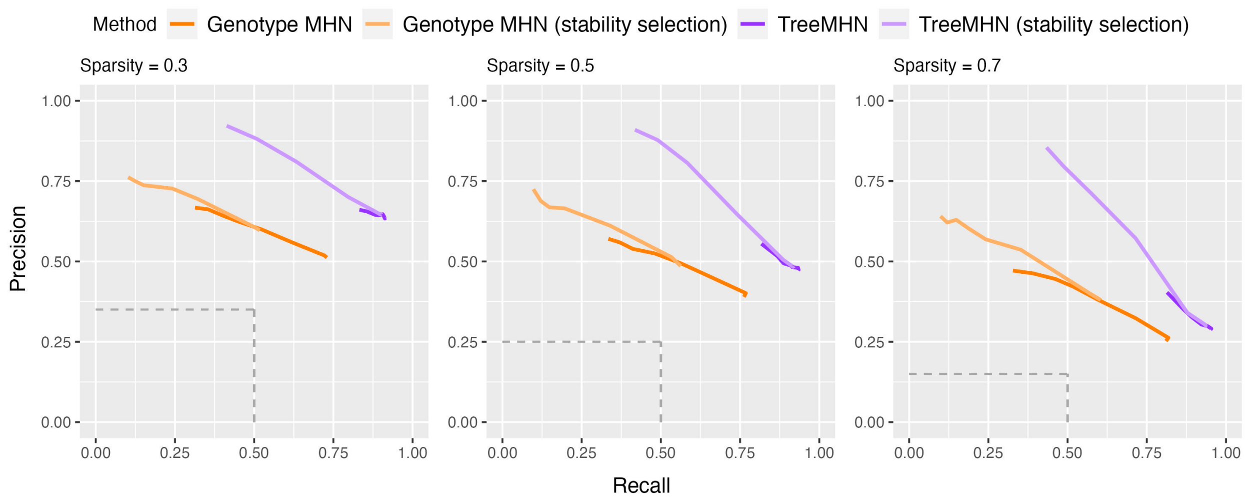

Figure S7: Performance of TreeMHN and the genotype MHN method (with and without stability selection) on simulated trees evaluated at different sparsity levels of the networks for $n = 10$ and $N = 500$. A sparsity level of 0.5 means that 50% of the off-diagonal entries in $\Theta$ are zero. The precision of random guess decreases as network sparsity increases.

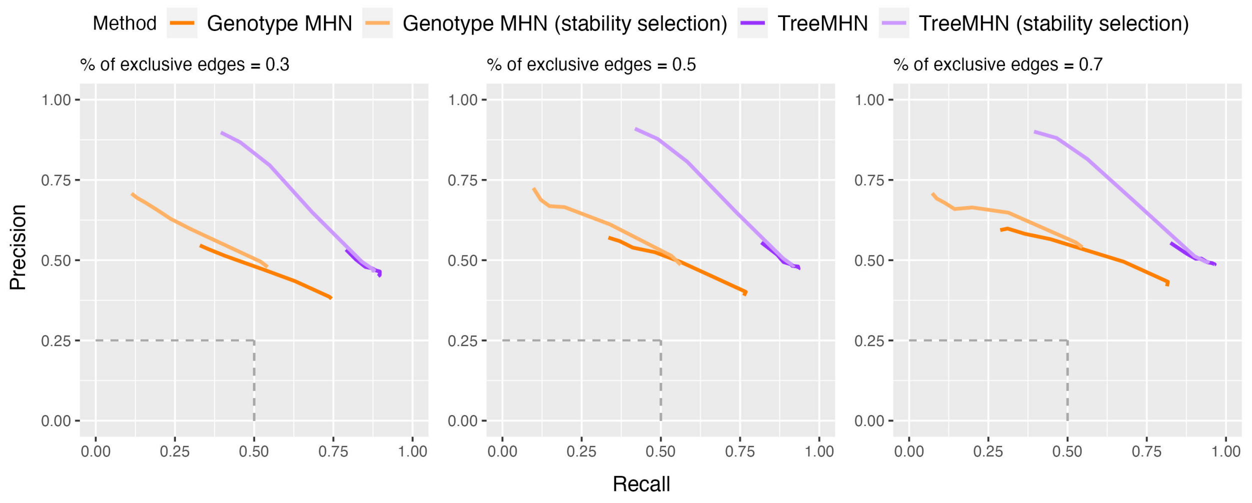

Figure S8: Performance of TreeMHN and the genotype MHN method (with and without stability selection) on simulated trees evaluated at different percentages of exclusive edges in $\Theta$ for $n = 10$ and $N = 500$. A percentage of exclusive edges equal to 0.5 means that 50% of the non-zero off-diagonal entries in $\Theta$ are negative.

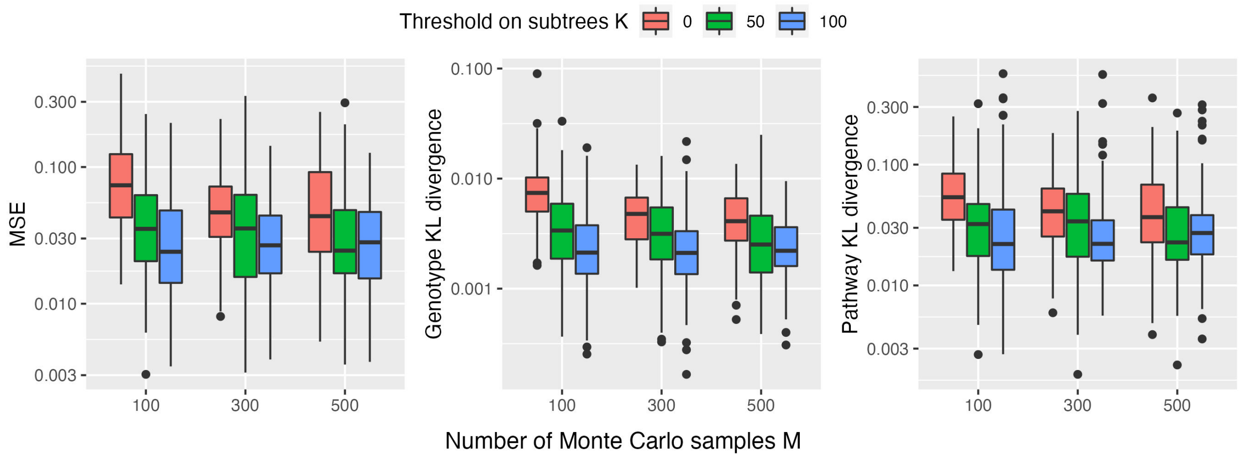

Figure S9: Comparison between the TreeMHN networks estimated using the MLE and the hybrid MC-EM methods for $n = 10$ and $N = 200$. For the hybrid MC-EM algorithm, we consider the thresholds on the subtrees $K \in \{0, 50, 100\}$ and the number of Monte Carlo samples $M \in \{100, 300, 500\}$. For each tree in the data set, we compute the expected time differences in exact form if its number of subtrees is below $K$. Otherwise, we use the approximation with importance sampling (Supplementary Section A.4). For each combination of $K$ and $M$, we run 100 simulations and evaluate the mean squared error (MSE), the genotype KL divergence [4], and the trajectory KL divergence (Methods) between the two output matrices. Overall, the higher the threshold $K$ and the more samples $M$, the smaller the differences among the networks estimated using the two inference methods.

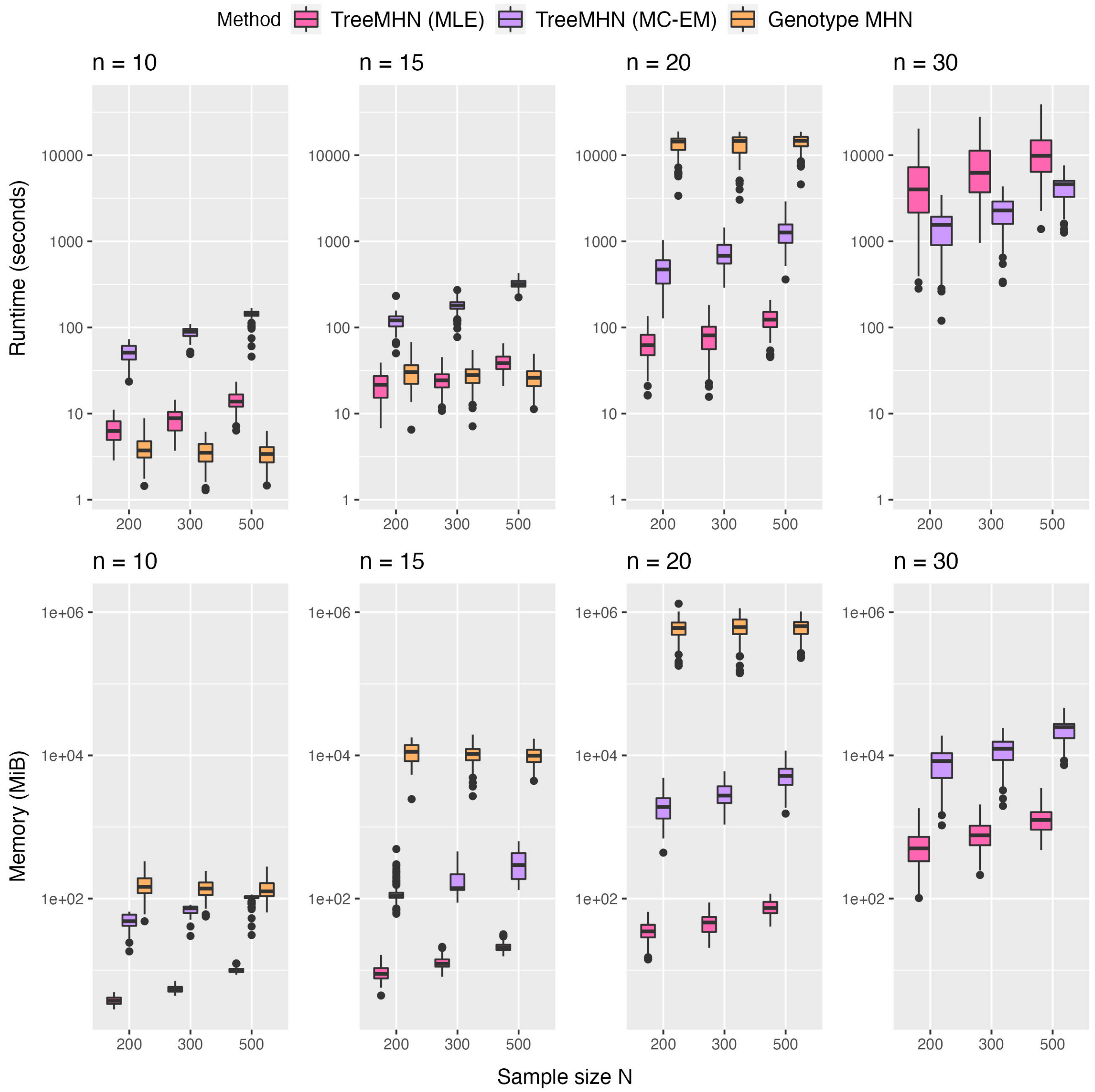

Figure S10: (a) Runtimes (wall-clock time in seconds) and (b) memory usage (in MiB) of TreeMHN (MLE and MC-EM) and the genotype MHN over 100 simulation runs for $n \in \{10, 15, 20, 30\}$, $N \in \{200, 300, 500\}$, and $\gamma = 1$. For the MC-EM version of TreeMHN, the threshold on the number of subtrees is $K = 50$, and the number of Monte Carlo samples is $M = 300$ for all configurations and simulation runs. For each method, each simulation run was performed on one CPU core of the AMD EPYC 7H12 processor on the ETH Euler cluster. The space and time complexity of the genotype MHN method increase exponentially in the number of mutations, so it is excluded in the $n = 30$ case. The runtime and memory usage of TreeMHN increase with the sample size, because the algorithm needs to go through every edge in the trees (Supplementary Section A.4). On the contrary, the genotype MHN method always summarize the genotypes into a vector of length $2^n$, so larger sample size may even improve the convergence of the optimization problem. Since we did not run REVOLVER and HINTRA using the original code but rather computed their key matrices (Methods), the runtime and memory usage of these two methods are not directly comparable. We use the R package `profmem` for memory profiling [7].

### E Application to acute myeloid leukemia data

#### E.1 Preprocessing of the data

The dataset we analyze is a cohort of $N = 123$ AML patient samples, which contains 543 somatic mutations in $n = 31$ cancer-associated genes [8]. Each mutation tree, reconstructed by SCITE [9], corresponds to the complete evolutionary history of a tumor. SCITE assumes the infinite sites assumption for individual genomic bases. We summarize the point mutations at the gene level such that the same gene may appear in parallel lineages of a tree. Also, there may be repeated mutations along the same lineage ($A \to B \to \cdots \to B$) due to gene-level summary, so we merge all such mutations to the position of their first occurrence. This rule also applies to the same mutations originating from a common direct ancestor ($B \leftarrow A \to B$).

#### E.2 Additional figures

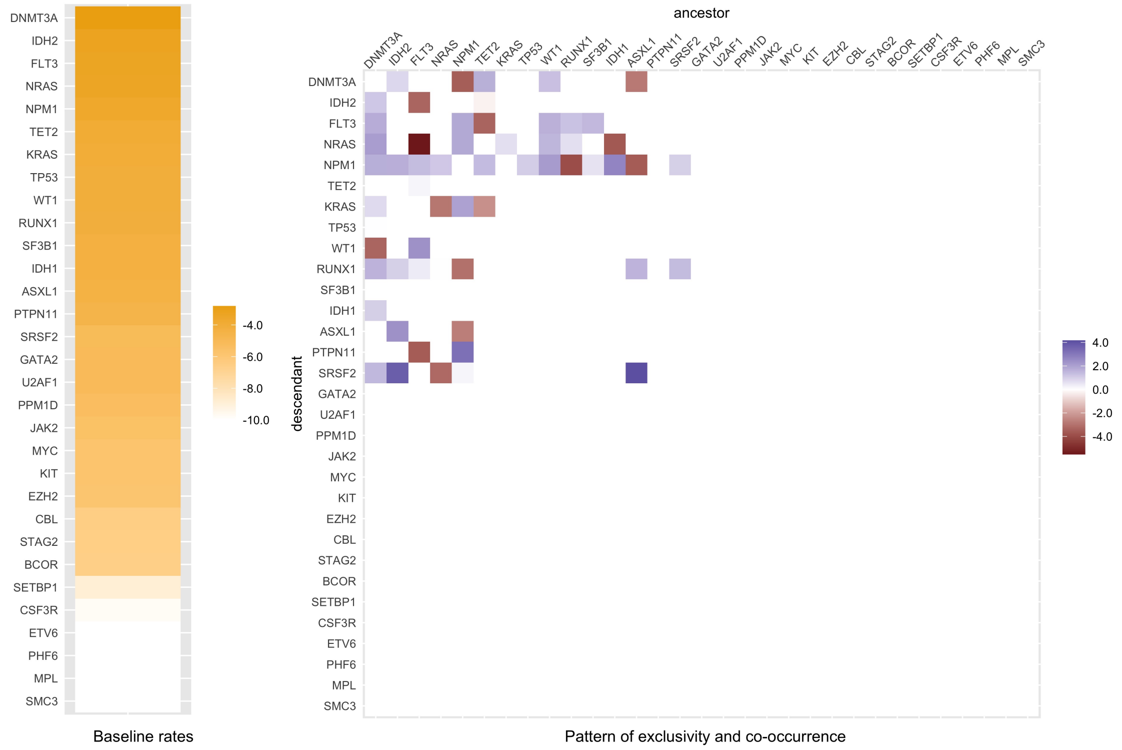

Figure S11: Full Mutual Hazard Network learned from a cohort of 123 AML patient samples [8]. This network is learned by following the stability selection procedure (Supplementary Section B) with threshold $\delta = 95\%$ and penalization parameter $\gamma = 0.1$. On the left it shows the baseline rates of the 31 mutational events, whereas the pairwise exclusive and co-occurring effects are illustrated on the right. Note that the values are displayed in logarithmic scale.

| | Trajectory | Probability | Count |
|---|---|---|---|
| 1 | Root->DNMT3A | 0.086 | 46 |
| 2 | Root->NRAS | 0.0755 | 45 |
| 3 | Root->FLT3 | 0.0721 | 47 |
| 4 | Root->IDH2 | 0.0601 | 33 |
| 5 | Root->NPM1 | 0.0402 | 49 |
| 6 | Root->KRAS | 0.0398 | 19 |
| 7 | Root->TET2 | 0.0371 | 20 |
| 8 | Root->TP53 | 0.0364 | 14 |
| 9 | Root->RUNX1 | 0.0325 | 25 |
| 10 | Root->SF3B1 | 0.0272 | 11 |
| 11 | Root->IDH1 | 0.0238 | 15 |
| 12 | Root->WT1 | 0.0238 | 18 |
| 13 | Root->PTPN11 | 0.0226 | 18 |
| 14 | Root->ASXL1 | 0.0209 | 17 |
| 15 | Root->SRSF2 | 0.0129 | 25 |
| 16 | Root->DNMT3A->NRAS | 0.0121 | 18 |
| 17 | Root->DNMT3A->FLT3 | 0.0096 | 14 |
| 18 | Root->IDH2->SRSF2 | 0.0081 | 12 |
| 19 | Root->NPM1->FLT3 | 0.0066 | 14 |
| 20 | Root->NPM1->PTPN11 | 0.0061 | 7 |
| 21 | Root->FLT3->WT1 | 0.0057 | 6 |
| 22 | Root->FLT3->NPM1 | 0.0056 | 9 |
| 23 | Root->ASXL1->SRSF2 | 0.0056 | 7 |
| 24 | Root->IDH1->NPM1 | 0.0055 | 5 |
| 25 | Root->NPM1->NRAS | 0.0054 | 10 |
| 26 | Root->TET2->DNMT3A | 0.0054 | 7 |
| 27 | Root->FLT3->WT1->NPM1 | 0.0043 | 2 |
| 28 | Root->NRAS->NPM1 | 0.004 | 6 |
| 29 | Root->IDH2->NPM1 | 0.004 | 8 |
| 30 | Root->DNMT3A->FLT3->NPM1 | 0.0039 | 2 |
| 31 | Root->RUNX1->FLT3 | 0.0031 | 4 |
| 32 | Root->SF3B1->FLT3 | 0.0028 | 5 |
| 33 | Root->NPM1->KRAS | 0.0027 | 6 |
| 34 | Root->DNMT3A->IDH2 | 0.0027 | 8 |
| 35 | Root->TET2->NPM1 | 0.0026 | 8 |
| 36 | Root->DNMT3A->RUNX1 | 0.0026 | 6 |
| 37 | Root->DNMT3A->NPM1 | 0.0026 | 20 |
| 38 | Root->DNMT3A->NPM1->FLT3 | 0.0024 | 6 |
| 39 | Root->NRAS->DNMT3A | 0.0023 | 1 |
| 40 | Root->FLT3->DNMT3A | 0.0023 | 2 |
| | | Median | 10.5 |

(a)

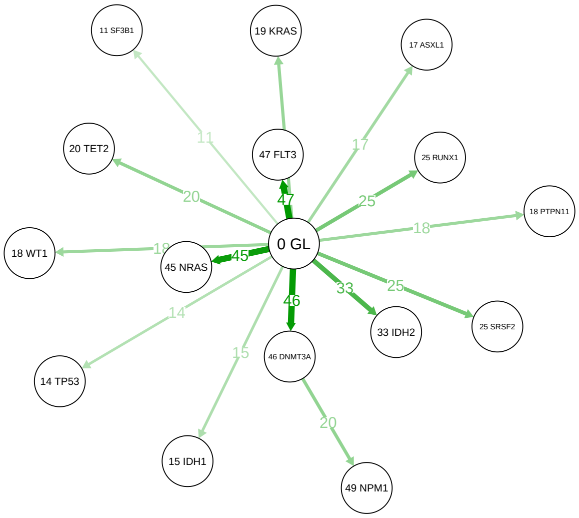

(b)

Figure S12: Comparison between TreeMHN and CONETT [10] on the inferred evolutionary trajectories from the AML dataset. (a) Table of top 40 most probable evolutionary trajectories inferred by TreeMHN. Each row corresponds to a trajectory along with its inferred probability and the number of times it appears in the transitive closures of the mutation trees (*i.e.* if there are directed edges from A to B and from B to C, then we add an edge from A to C). (b) Germline-rooted evolutionary trajectory tree computed by CONETT, which is conserved in at least 10 patients at 10% significance level. The number on an edge indicates the number of patients having the path from the GL node to the node which the edge is pointing to.

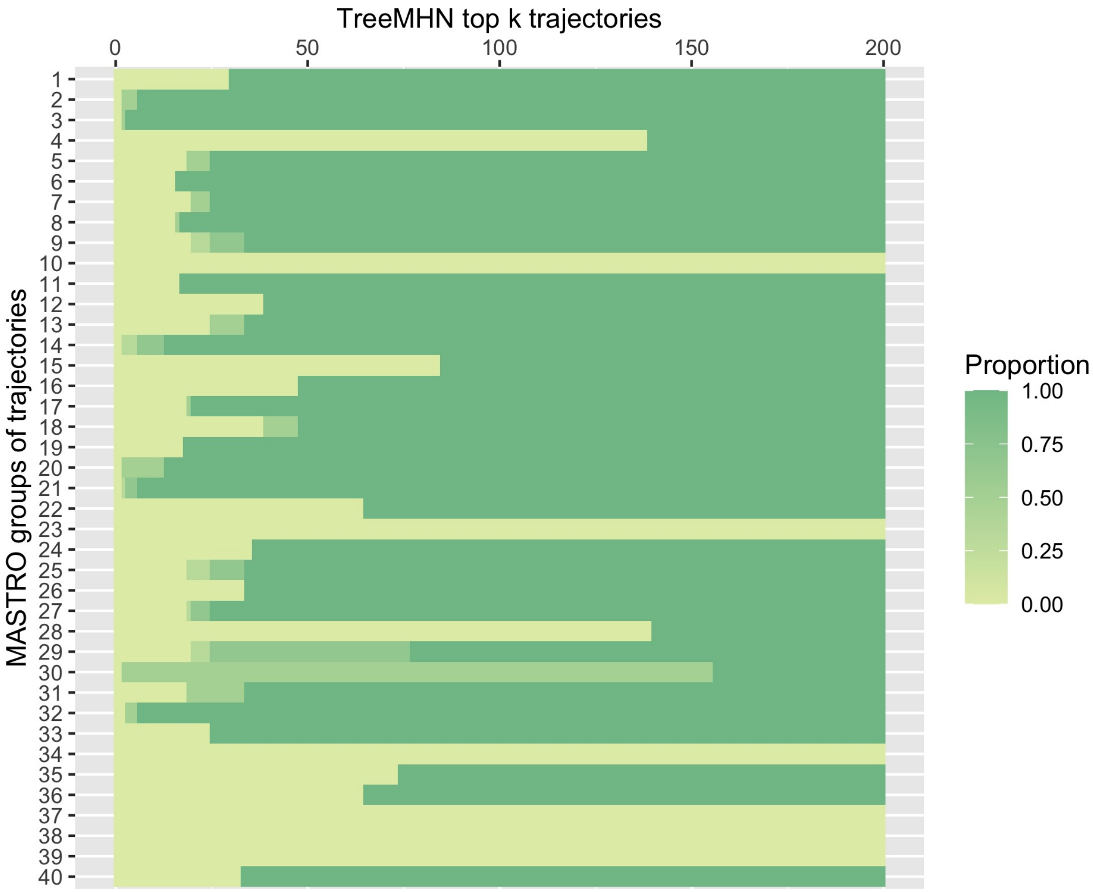

Figure S13: Proportion of significantly conserved trajectories reported by MASTRO [11] found in the top $k$ most probable trajectories predicted by TreeMHN for the AML dataset. Here we consider $k \leq 200$. Each row corresponds to one group in Supplementary Figure 5 of [11].

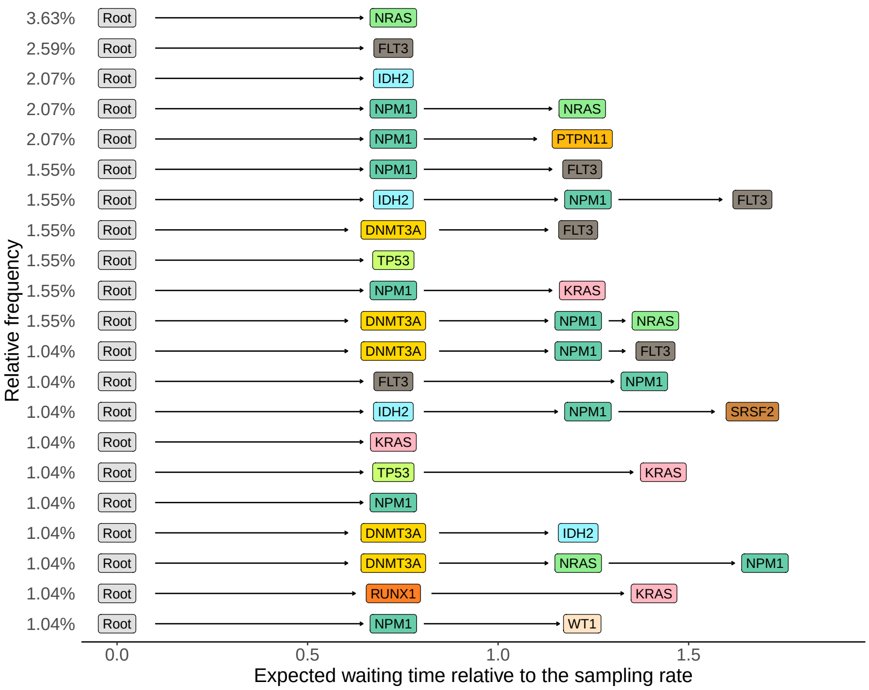

Figure S14: Observed evolutionary trajectories that appear at least twice in the AML tumor trees [8]. The rows represent the evolutionary trajectories, labeled and ordered by their relative frequencies. The horizontal positions of the mutations correspond to their expected waiting times relative to the sampling rate of $\lambda_s = 1$, which are computed based on the estimated AML network. Despite the small sample size, the relative frequencies of the trajectories still match closely with the estimated trajectory probabilities.

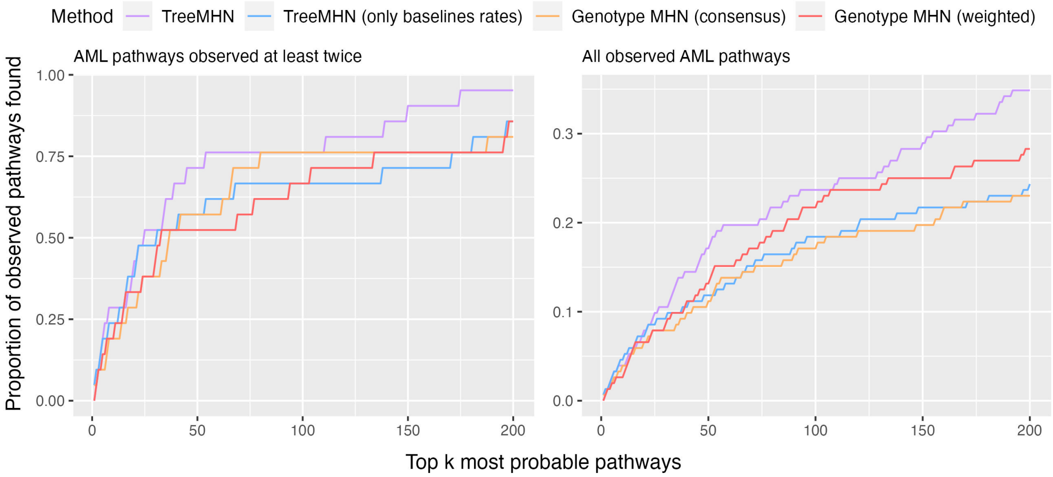

Figure S15: Proportion of observed AML trajectories found in the top $k$ most probable trajectories predicted by four methods: TreeMHN, TreeMHN with only baseline rates, genotype MHN using the consensus subclonal genotypes, genotype MHN using the weighted subclonal genotypes (Supplementary Section C.2). Here we consider $k \leq 200$. On the left we show the comparison on the AML trajectories observed at least twice in the cohort, whereas the one on the right includes all observed trajectories. For scalability, we run both variants of genotype MHNs by focusing on the top 15 mutations that have non-zero off-diagonal entries in the estimated network (Supplementary Figure S11).

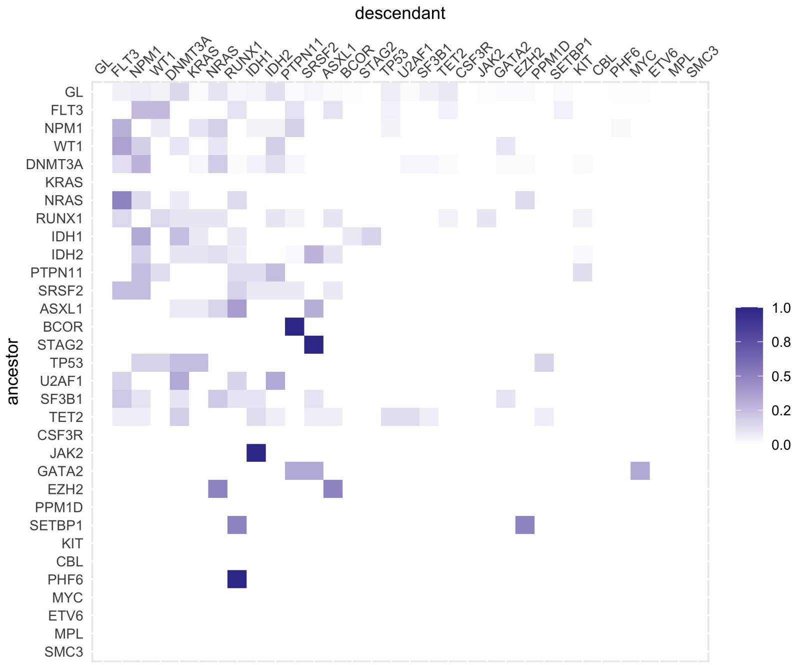

Figure S16: REVOLVER's information transfer matrix $\boldsymbol{w}$ estimated on the AML dataset after normalization. Each entry $w_{ij}$ represents the empirical probability of mutation $j$ being the descendant of mutation $i$.

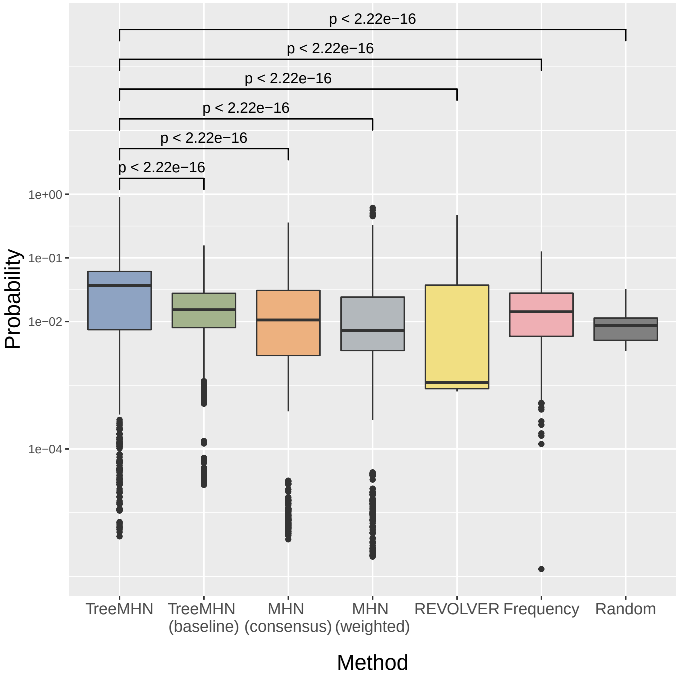

Figure S17: Performance assessment on retrospective predictions on the AML dataset. Each point in the plot corresponds to the probability of a downstream event given a rooted subtree of a primary tumor tree. The last column shows the probabilities of random guess, where each possible event gets equal weight. Since for each rooted subtree, the number of possible events is different depending on the size of the tree, the last boxplot is not a line. The p-values for the one-sided Wilcoxon signed-rank tests between TreeMHN and alternative methods are also displayed.

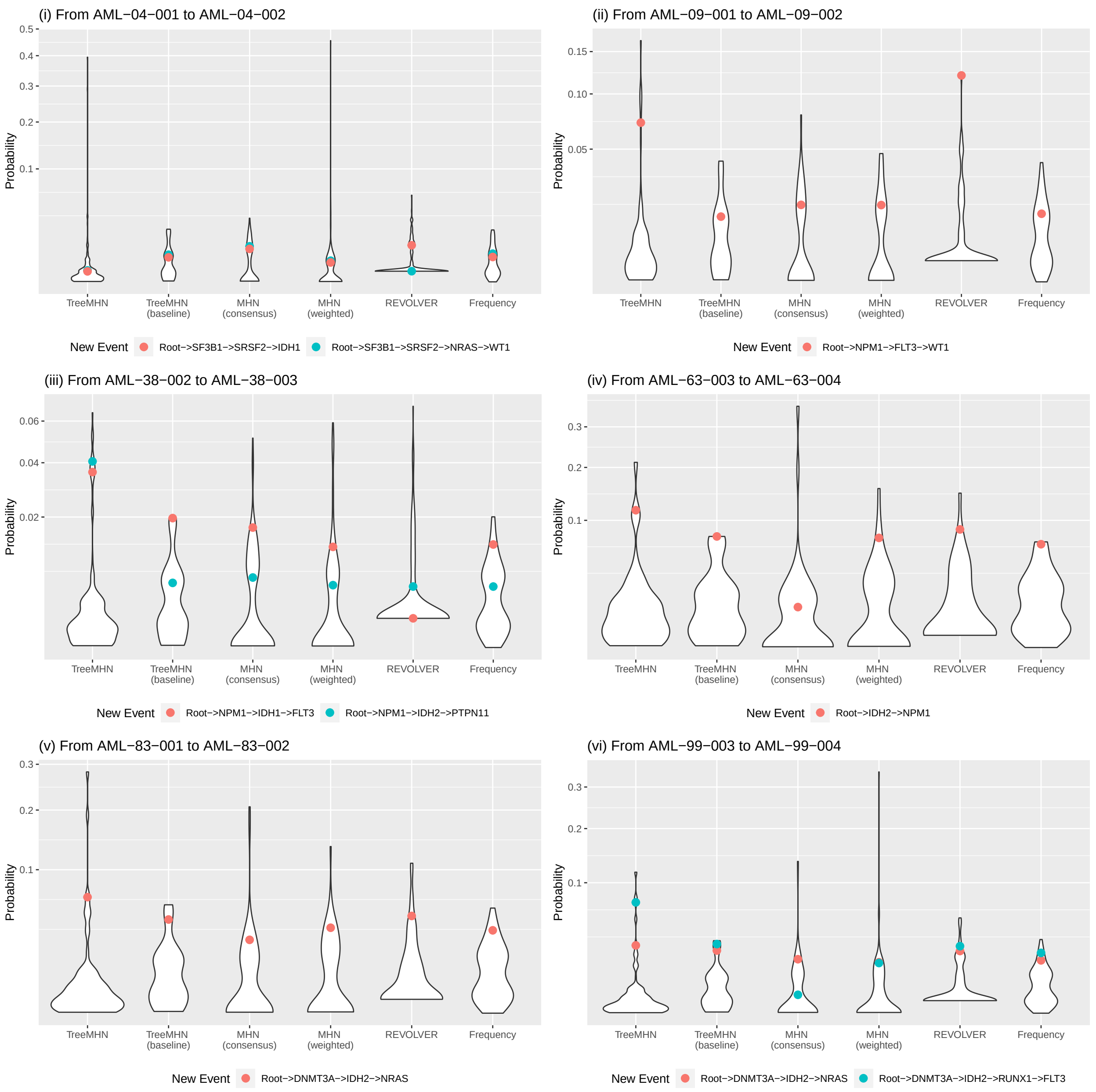

Figure S18: Performance assessment on forward predictions on the AML longitudinal samples. Each panel corresponds to a pair of consecutive trees where the second tree has at least one new event. For each method, the violin plot shows the estimated probability distribution of all possible events, and the new events are colored dots. These events correspond to the events shown in Figure 6c of the main text.

### F Application to non-small-cell lung cancer data

#### F.1 Preprocessing of the data

To apply TreeMHN to the TRACERx NSCLC data set [12], we retrieve the uncorrelated phylogenetic trees for $N = 99$ patients and $n = 79$ driver mutations from the R package `evoverse.datasets` [13]. Each patient sample has multiple clone trees with different scores, and each node in the clone tree can have multiple mutations, whose ordering cannot be determined. Hence, for each clone tree, we enumerate or randomly sample the permutations of the mutations in the clones (depending on the clone size), and assign equal weights to the resulting fully resolved trees. Then, we multiply the weights by the score of the original clone tree. Finally, we use a weighted version of TreeMHN, where the total weights of the trees for each patient sum up to 1.

#### F.2 Additional figures

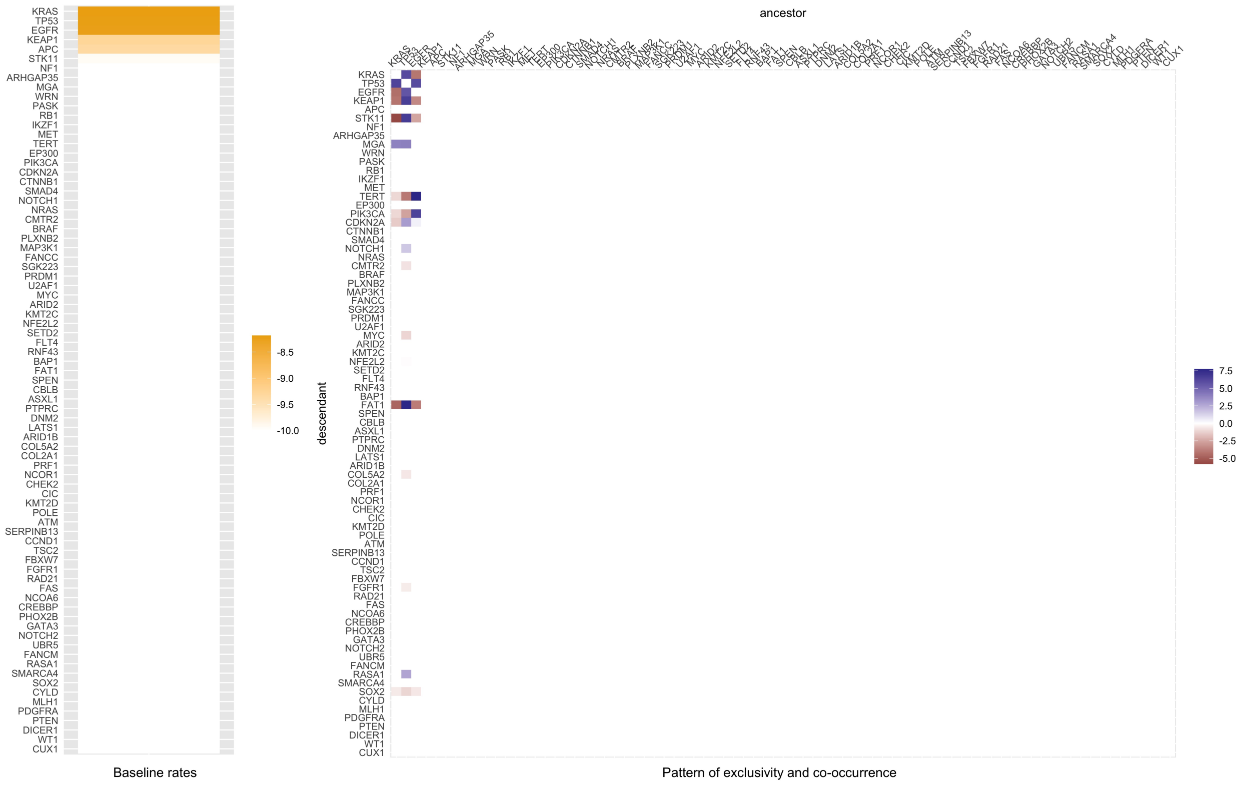

Figure S19: Full Mutual Hazard Network learned from a cohort of 99 NSCLC patient samples [5, 12]. This network is learned by following the stability selection procedure (Supplementary Section B) with threshold $\delta = 99\%$ and penalization parameter $\gamma = 0.001$. On the left it shows the baseline rates of the 79 mutational events, whereas the pairwise exclusive and co-occurring effects are illustrated on the right. Note that the values are displayed in logarithmic scale.

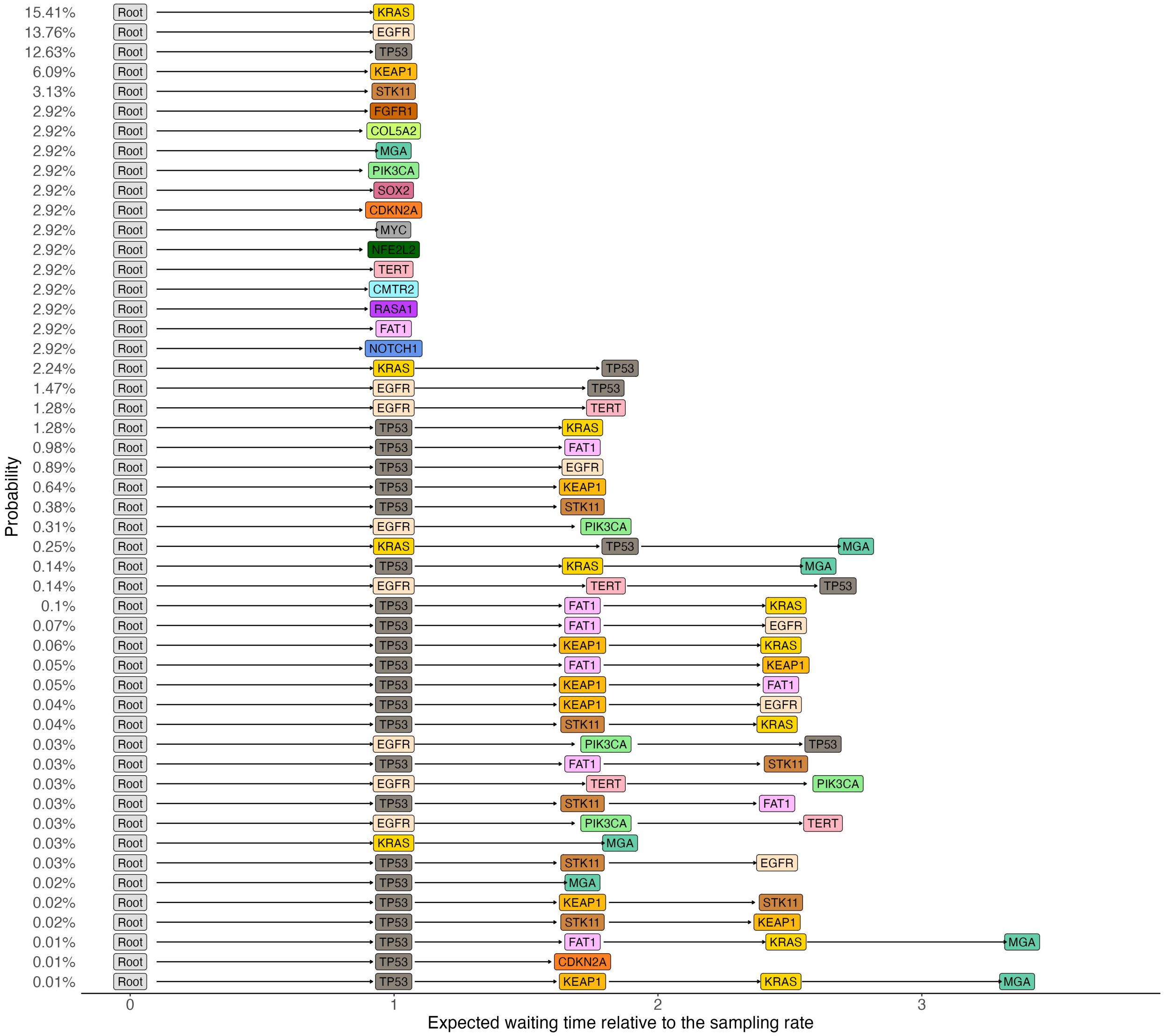

Figure S20: Top 50 most probable evolutionary trajectories inferred from the partial Mutual Hazard Network of the TRACERx NSCLC dataset (Figure 8). Note that the total number of evolutionary trajectories is in the order of 18!.

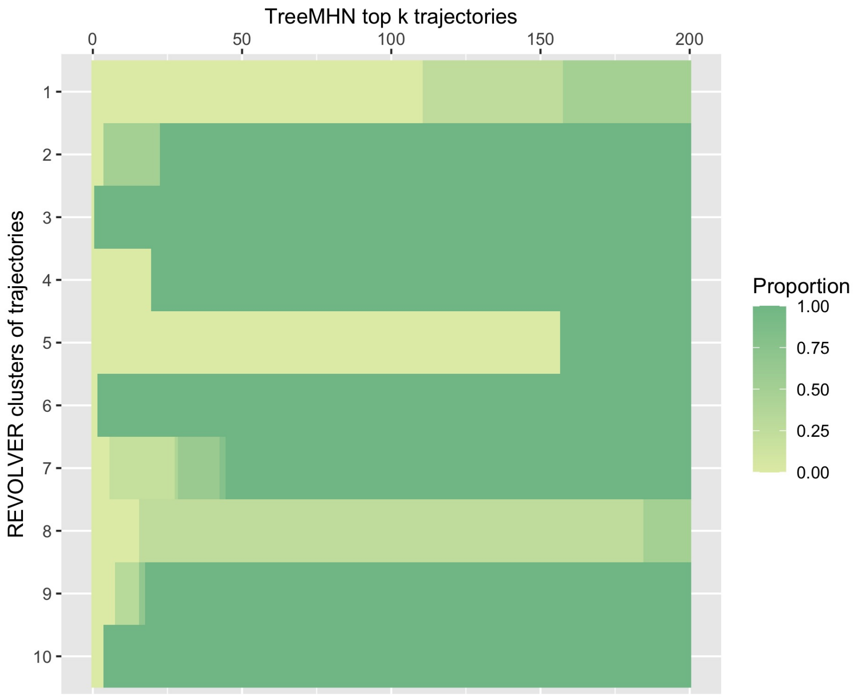

Figure S21: Proportion of repeated trajectories reported by REVOLVER found in the top $k$ most probable trajectories predicted by TreeMHN for the NSCLC dataset. Here we consider $k \leq 200$. Each line corresponds to one cluster in Supplementary Figure 7 of [5].

### G Application to breast cancer data

#### G.1 Preprocessing of the data

Following [14], we use the phylogenetic trees inferred by SPRUCE [15] for the breast cancer data in [16]. The number of patients in the source data is $N = 1756$. Christensen et al. [14] reduce this number to 1315 by focusing on SNVs in copy-neutral autosomal regions. We partition the patients into 8 subgroups according to their hormone receptor status (HR+/HER2+, HR+/HER2-, HR-/HER2+, Triple Negative) and sample type (primary vs. metastasis). For primary tumors, we consider only the treatment-free samples to avoid confounding signals. Moreover, we restrict the analysis to the union of mutations that appear in at least 10% of the patients in each subgroup. These preprocessing steps result in $n = 19$ mutations and $N = 1152$ patients with 1232 phylogenetic trees. Trees for the same patient sample get equal weights. The distributions of patients and mutations across subgroups are provided in Supplementary Table S1 and Supplementary Table S2 respectively.

| | HR+/HER2+ | HR+/HER2- | HR-/HER2+ | Triple Negative |
|---|---|---|---|---|
| Primary | 31 | 442 | 14 | 57 |
| Metastasis | 73 | 444 | 26 | 65 |

Table S1: Number of advanced breast cancer patients in each of the eight subgroup after the preprocessing steps.

| | HR+/HER2+ | HR+/HER2- | HR-/HER2+ | Triple Negative |
|---|---|---|---|---|
| Primary | 10 | 17 | 8 | 14 |
| Metastasis | 15 | 19 | 12 | 12 |

Table S2: Number of mutations in each of the eight subgroup after the preprocessing steps.

#### G.2 Additional figures

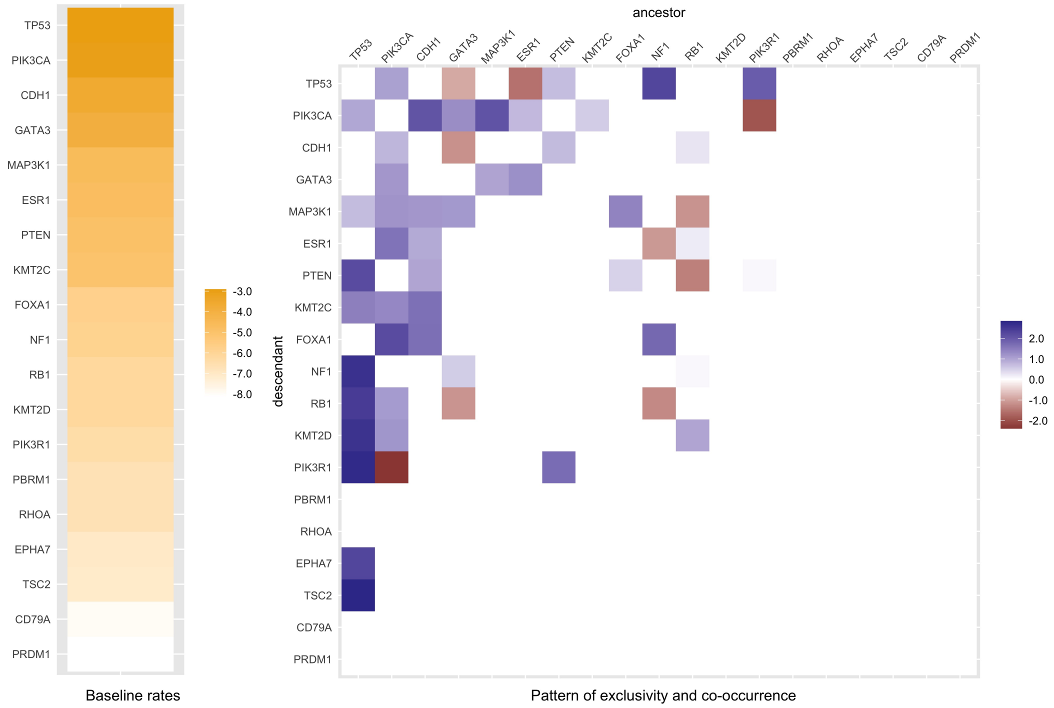

Figure S22: Full Mutual Hazard Network learned from a cohort of breast cancer patient samples [14, 16]. This network is learned by following the stability selection procedure (Supplementary Section B) with threshold $\delta = 99\%$ and penalization parameter $\gamma = 0.01$. On the left it shows the baseline rates of the 19 mutational events, whereas the pairwise exclusive and co-occurring effects are illustrated on the right. Note that the values are displayed in logarithmic scale.

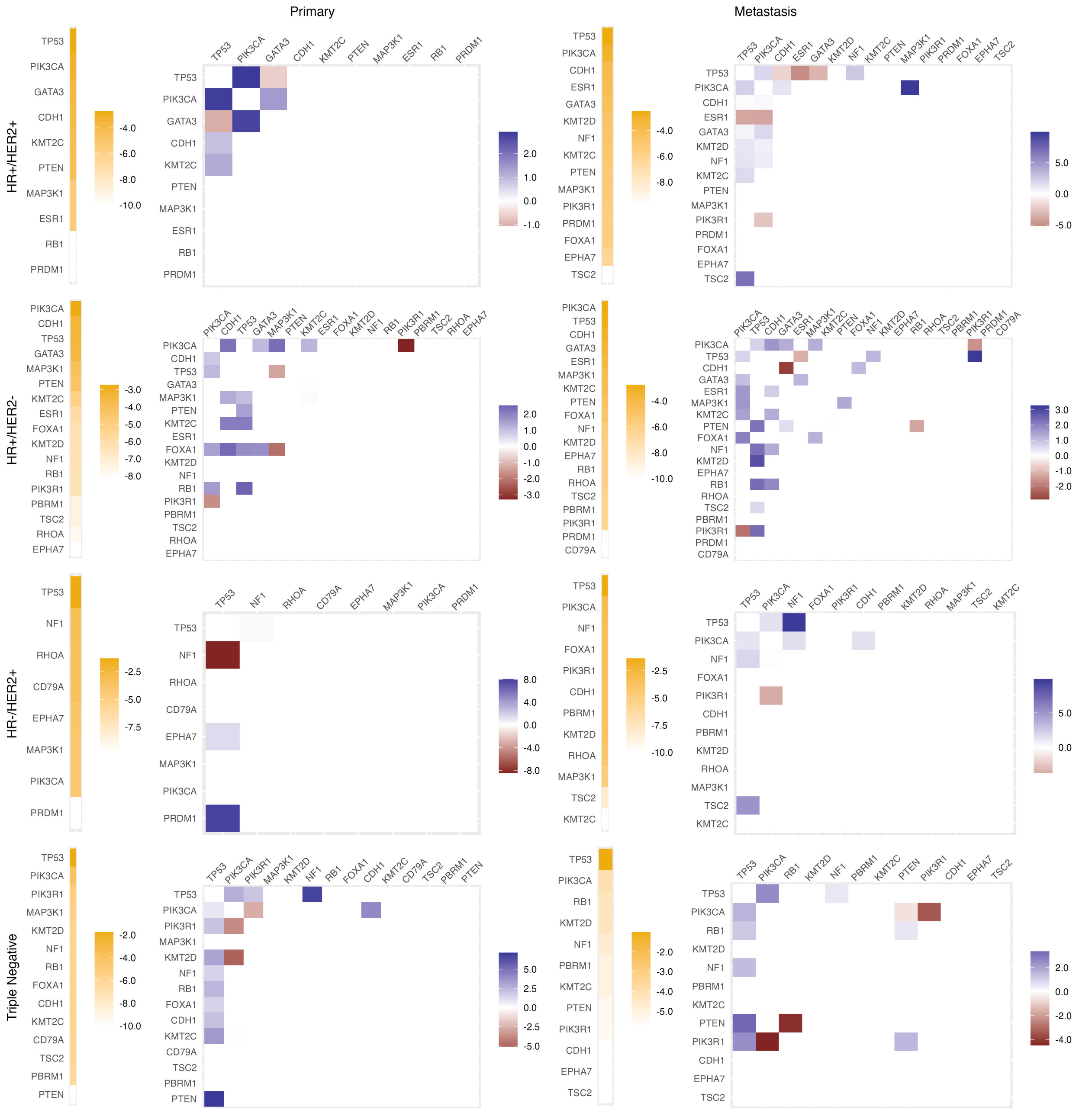

Figure S23: Full Mutual Hazard Networks for the eight subgroups defined in Supplementary Section G.1. Using the stability selection procedure with threshold $\delta = 99\%$ (Supplementary Section B), we control the expected false positive rate to be at most 10% by varying the penalty level $\gamma$ for each subgroup based on the number of mutations (Supplementary Table S2).

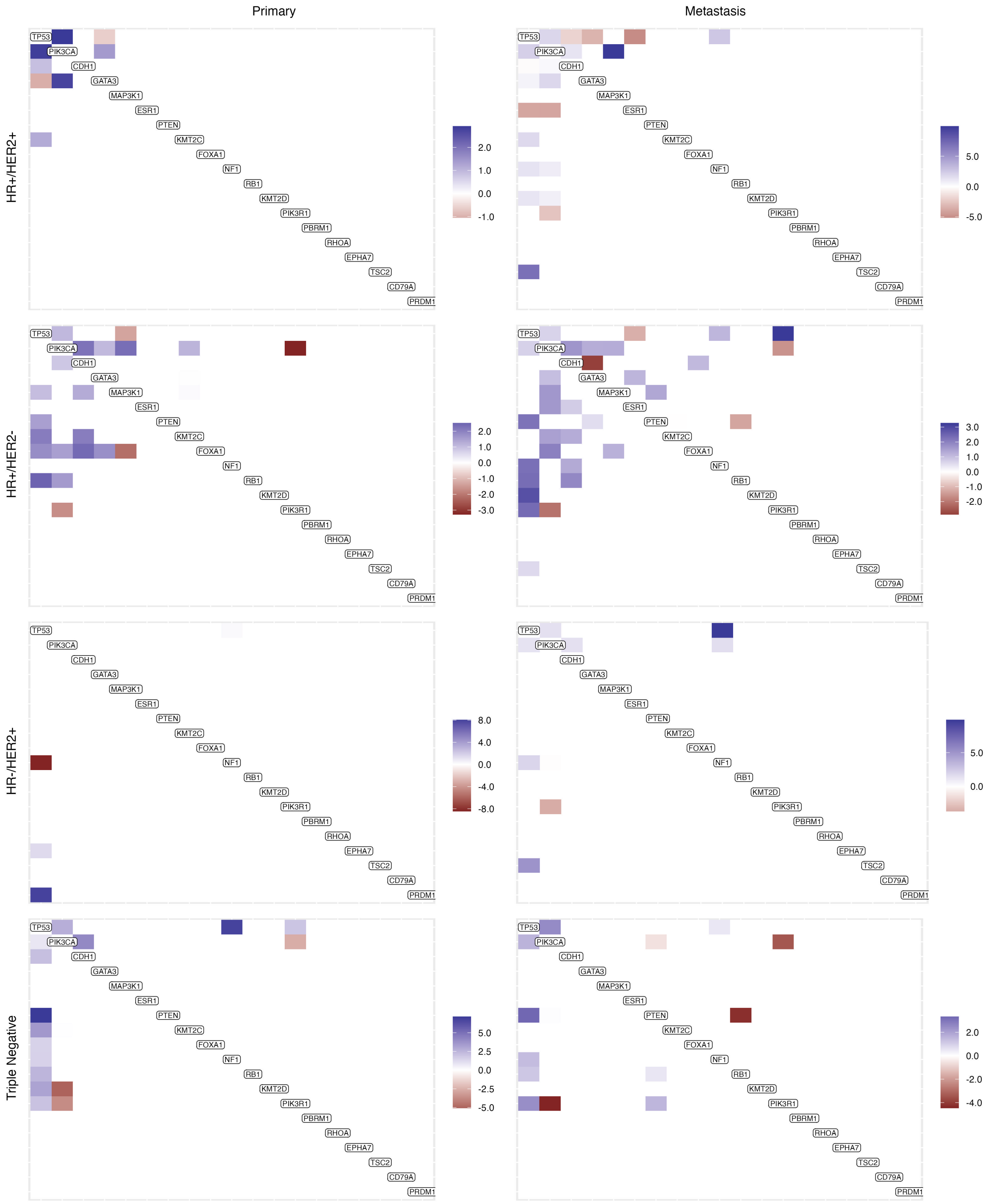

Figure S24: Partial Mutual Hazard Networks for the eight subgroups defined in Supplementary Section G.1. The columns and rows of these matrices are ordered by decreasing baseline rates in the main matrix (Supplementary Figure S22).

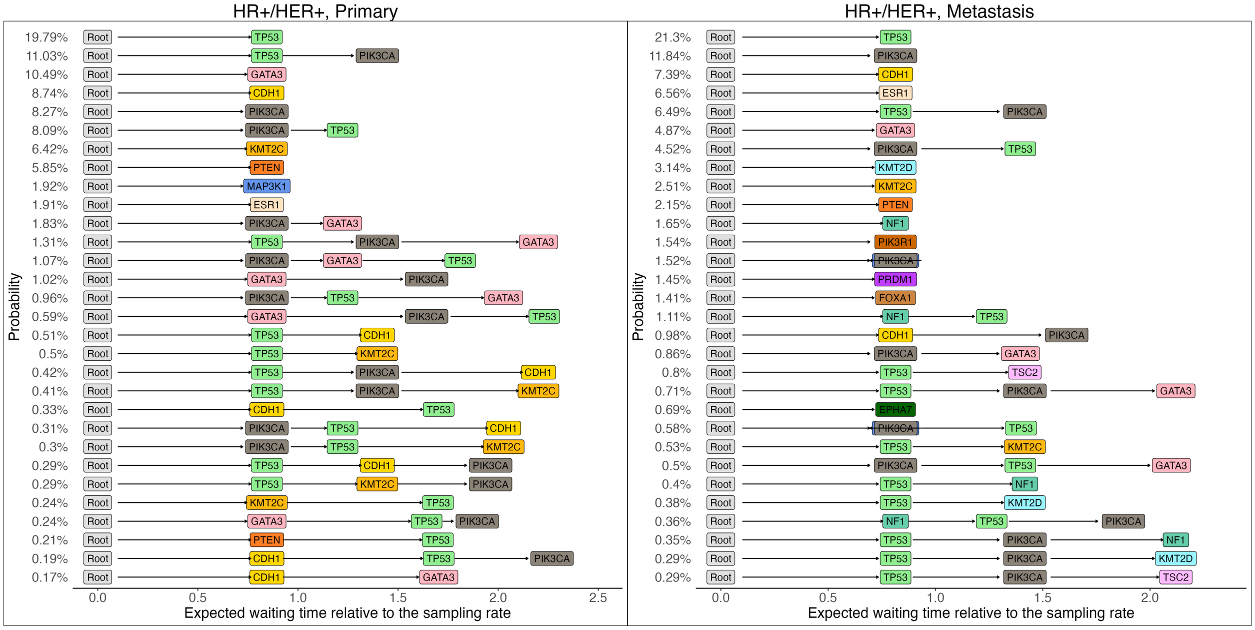

Figure S25: Top 30 most probable evolutionary trajectories of the HR+/HER+ primary (left panel) and metastasis (right panel) subgroups for the breast cancer data [16].

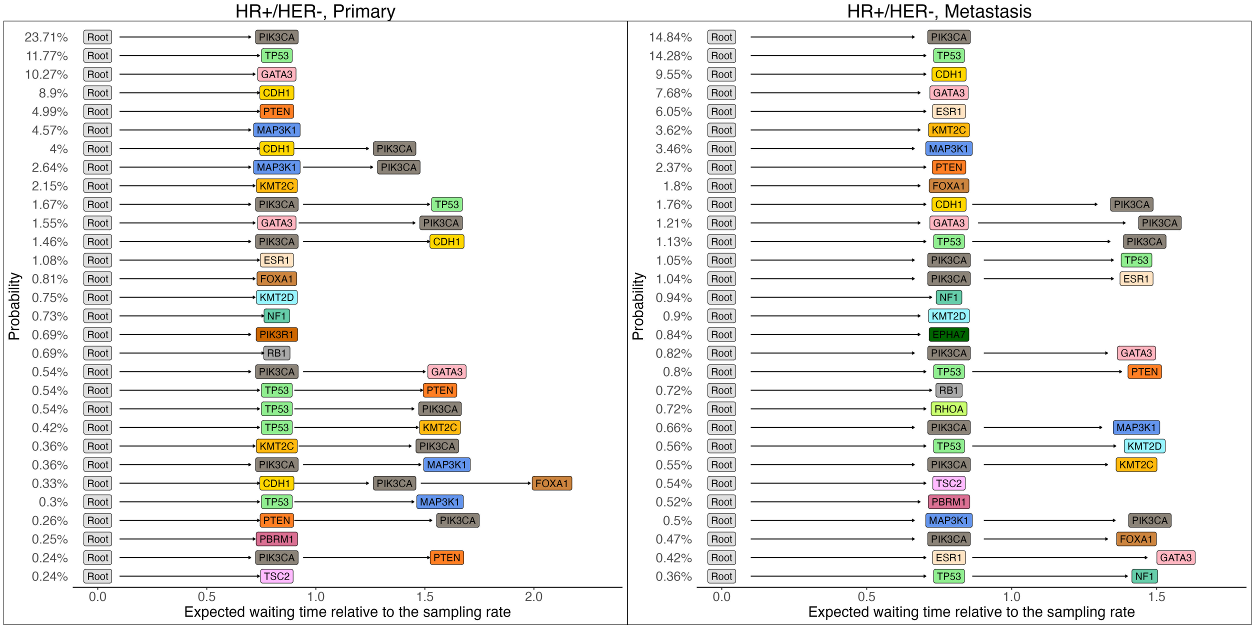

Figure S26: Top 30 most probable evolutionary trajectories of the HR+/HER- primary (left panel) and metastasis (right panel) subgroups for the breast cancer data [16].

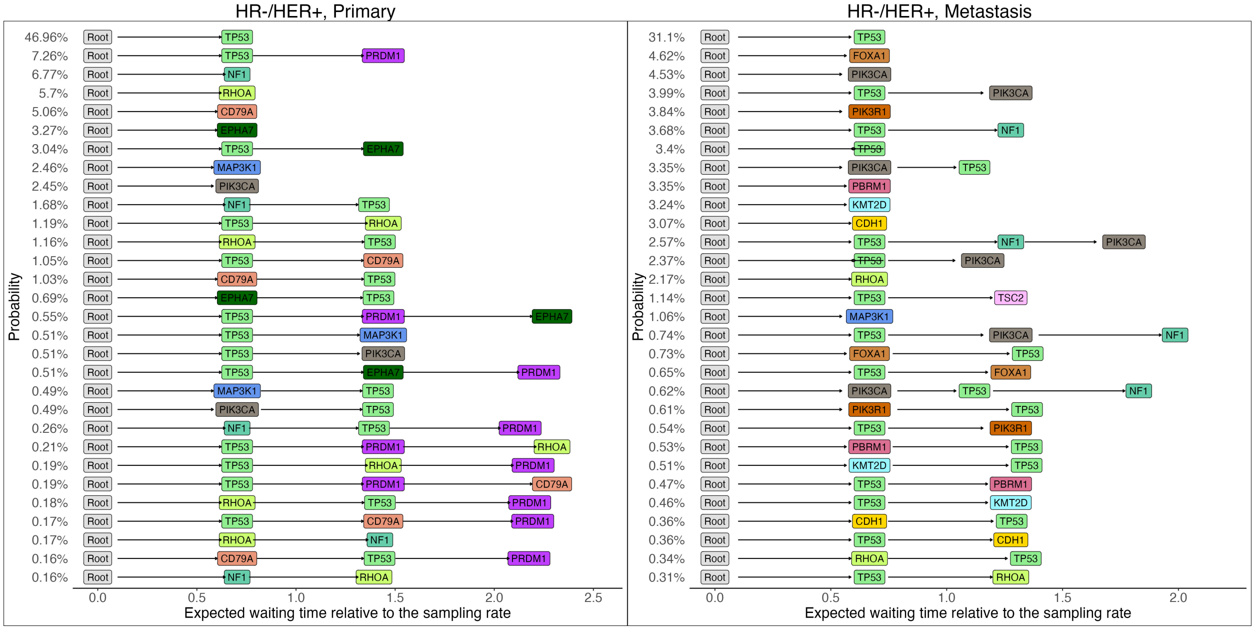

Figure S27: Top 30 most probable evolutionary trajectories of the HR-/HER+ primary (left panel) and metastasis (right panel) subgroups for the breast cancer data [16].

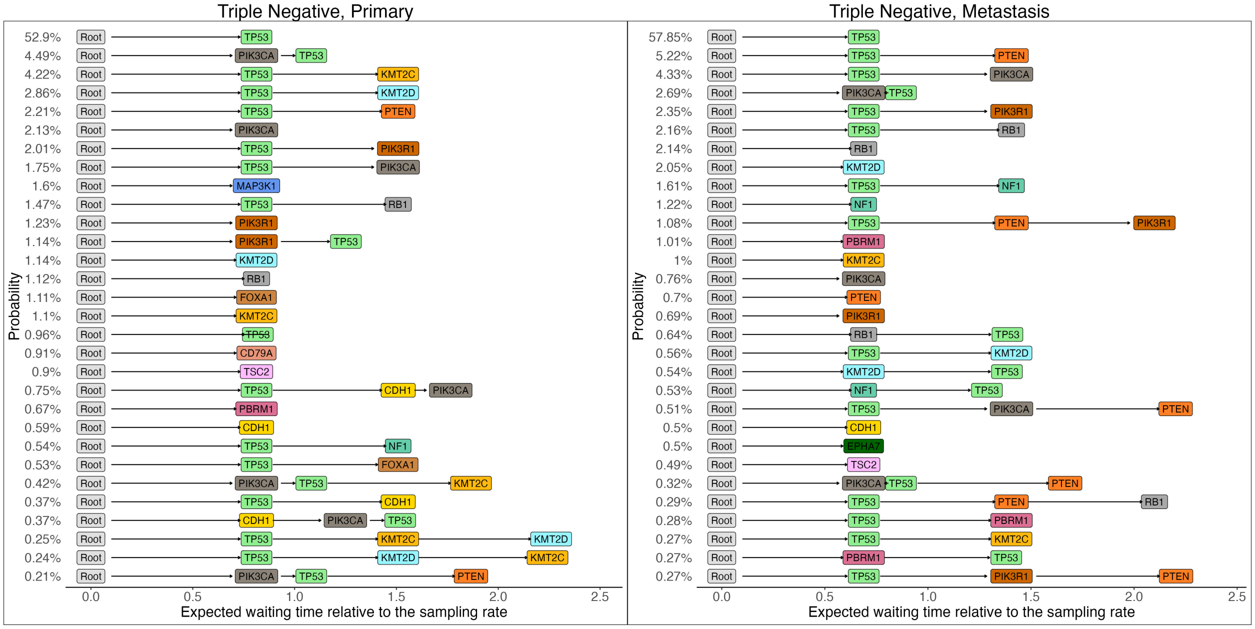

Figure S28: Top 30 most probable evolutionary trajectories of the triple-negative primary (left panel) and metastasis (right panel) subgroups for the breast cancer data [16].
